# Barrier Immune Memory is Promoted by Intestinal Epithelial Cell Presentation of Injected Bacterial Antigens

**DOI:** 10.64898/2026.03.27.714828

**Authors:** C. Garrett Wilson, M. Pragun Acharya, Laura Karsch, Lennard W. Duck, Nana Twumasi-Ankrah, Yuanyou Wang, Hongxing Shen, Chuan Xing, Blake F. Frey, Vishal Oza, Stacey N. Harbour, Yoshiko Nagaoka-Kamata, Jeffrey R. Singer, Robin D. Hatton, Jeffrey Moffitt, Matthias Gunzer, Carlene L. Zindl, Casey T. Weaver

**Affiliations:** Department of Pathology, Heersink School of Medicine, University of Alabama at Birmingham, AL, USA; Institute for Experimental Immunology and Imaging, University Hospital Essen, University Duisburg-Essen, Essen, Germany; Department of Medicine, Heersink School of Medicine, University of Alabama at Birmingham, AL, USA; Program in Cellular and Molecular Medicine, Boston Children’s Hospital, Boston, MA, USA; Medical Scientist Training Program, University of Alabama at Birmingham, AL, USA; Department of Medicine, Vanderbilt University Medical Center, Nashville, TN

## Abstract

The contributions of antigen compartmentalization to recognition differences between CD4 and CD8 T cells have long been appreciated, but little is known of how subcellular localization of different antigens expressed by a single pathogen impacts T cell immunity. By tracking a clonal CD4 T cell response to its cognate epitope shuttled between different virulence proteins of the enteropathogenic bacterium, *Citrobacter rodentium* (*Cr*), we find a remarkable bias in the magnitude and quality of the response contingent on whether antigen remains bacterially associated or is introduced into intestinal epithelial cells colonized by the bacterium. Only proteins injected into the cytosol of colonocytes via the type 3 secretion system (T3SS) of *Cr* were found to recruit robust antigen-specific T cell responses to the infected mucosa and give rise to CD4 resident memory T (Trm) cells that populate the mucosal epithelium—and this required direct presentation of these antigens by infected epithelial cells. Single-cell transcriptomic analyses revealed that sustained, bidirectional epithelial-T cell communication was required both to elicit epithelial barrier-protective T cell help and to promote transcriptional networks that program a tissue-residency rather than central memory fate. These results establish a central role for antigen presentation by non-professional APCs in controlling memory fate decisions by CD4 T cells, with important implications for development of successful mucosal vaccines.

## Introduction

The guiding principle of adaptive immunity is antigen recognition. For T cells, that recognition is restricted to the surface of cells that display antigenic peptides bound to major histocompatibility (MHC) molecules: MHCI for CD8 T cells and MHCII for CD4 T cells. Whereas MHCI is ubiquitously expressed by nucleated cells, expression of MHCII is more restricted, primarily limited to so-called “professional” antigen-presenting cells (APCs) of hematopoietic origin (i.e., dendritic cells, macrophages, and B cells). However, some non-hematopoietic cells—including epithelial cells lining barrier tissues at sites of pathogen entry^1–7^—can be induced to express MHCII in the context of an immune response and may act as “non-professional” APCs. Although these cells lack costimulatory molecules and cannot activate naive T cells^8–10^, they may recruit CD4 T cell help to combat infections via presentation of pathogen-derived antigens—akin to the recruitment of cytotoxic CD8 T cells by infected MHCI-positive cells to target them for death.

We recently identified a subset of absorptive intestinal epithelial cells (IECs) termed distal colonocytes (DCCs)—so named for their unique distribution to distal colon—that are targeted by the murine enteropathogenic bacterium, *Citrobacter rodentium* (*Cr*)^1^. *Cr* belongs to a family of closely related Gram-negative bacteria that includes the human pathogens enterohaemorrhagic and enteropathogenic *E. coli* (EHEC and EPEC)^11,12^, which require deployment of a type III secretion system (T3SS) to colonize the apical surface of host IECs, where they form characteristic attaching and effacing (A/E) lesions^11,15,16^. Bacterial virulence proteins encoded by genes of the locus of enterocyte effacement (LEE) pathogenicity island^13,14^ are expressed early in infection to assemble the T3SS molecular syringe that docks to the apical surface of IECs and injects bacterial effector proteins into the epithelial cell, where they localize to the apical cell membrane or cytosol^17–20^. Both LEE and non-LEE gene loci (e.g., *Nle* family) encode bacterial effectors that are secreted via the T3SS and contribute to *Cr* pathogenicity^21,22^. DCCs—but not absorptive colonic epithelial cells (cECs) that line the proximal colon (proximal colonocytes; PCCs)—were found to recruit CD4 T cells that produced the epithelium-protective cytokine IL-22 contingent on their expression of MHCII^1^, establishing a role for presentation by DCCs of *Cr* antigens loaded onto these cECs and reflecting the tropism of *Cr* for this epithelial population^1^. However, the range of bacterial virulence proteins that contribute to the CD4 T cell response and how their compartmentalization between the bacterium or host epithelial cell—and subcellular compartments therein—might affect the amplitude and quality of this response are undefined.

Using isogenic strains of *Cr* engineered to express the same CD4 T cell epitope fused to different *Cr* virulence proteins, we have mapped a clonal CD4 T-cell response based on the type of bacterial “carrier” protein used to deliver cognate antigen—enabling assessment of the T-cell response without the confounding variable of differing affinities of antigenic peptides for MHCII. We find striking effects on the magnitude and quality of the clonal CD4 T cell response contingent on bacterial versus epithelial cell antigen localization, with bacterial antigens that are injected into the cytosol of epithelial cells promoting the greatest clonal expansion and retention of T cells in the infected mucosa as well as their development into tissue-resident memory (Trm) cells. In absence of antigenic localization to the cEC cytosol, the CD4 T cell clonal response is blunted, and memory cell development largely defaults to the central memory (Tcm) compartment, generating few memory cells that populate the intestinal epithelium where they might more rapidly respond to pathogen upon reinfection. These results have profound implications for the development of successful mucosal vaccines and establish a central role for antigen presentation by non-professional APCs in controlling memory fate decisions by CD4 T cells.

## Results

### Injected bacterial antigens drive enhanced mucosal CD4 T cell responses

To define how antigen localization of bacterial antigens influences CD4 T cell responses, we engineered *C. rodentium* strains that express the LCMV gp66-80 peptide recognized by the SMARTA T-cell receptor (TCR) fused to different bacterial proteins, including a hemagglutinin (HA) tag for tracking fusion protein expression^23^ (**Fig. 1a,b; ED Fig. 1a-c; and Supplementary Table 1**). Four bacterial proteins targeted to different bacterial or host cell compartments were selected: intimin (eae), a LEE-encoded surface-exposed adhesin that localizes to the outer bacterial cell membrane; the translocated intimin receptor (tir), a LEE-encoded protein that localizes to the apical plasma membrane following injection into cECs, where it binds intimin to establish firm bacterial-cEC adhesion; nleA, a non-LEE encoded effector that distributes diffusely throughout the host cell cytosol; and espZ, a LEE-encoded effector that concentrates in the cytosol beneath adherent bacteria (**Fig. 1b,d**). All engineered strains maintain virulence comparable to the wild-type, parent *Cr* strain (DBS100) in susceptible C3H/HeJ mice, confirming that epitope insertion did not compromise bacterial fitness or pathogenicity (**Fig. 1c**).

**Fig. 1:**
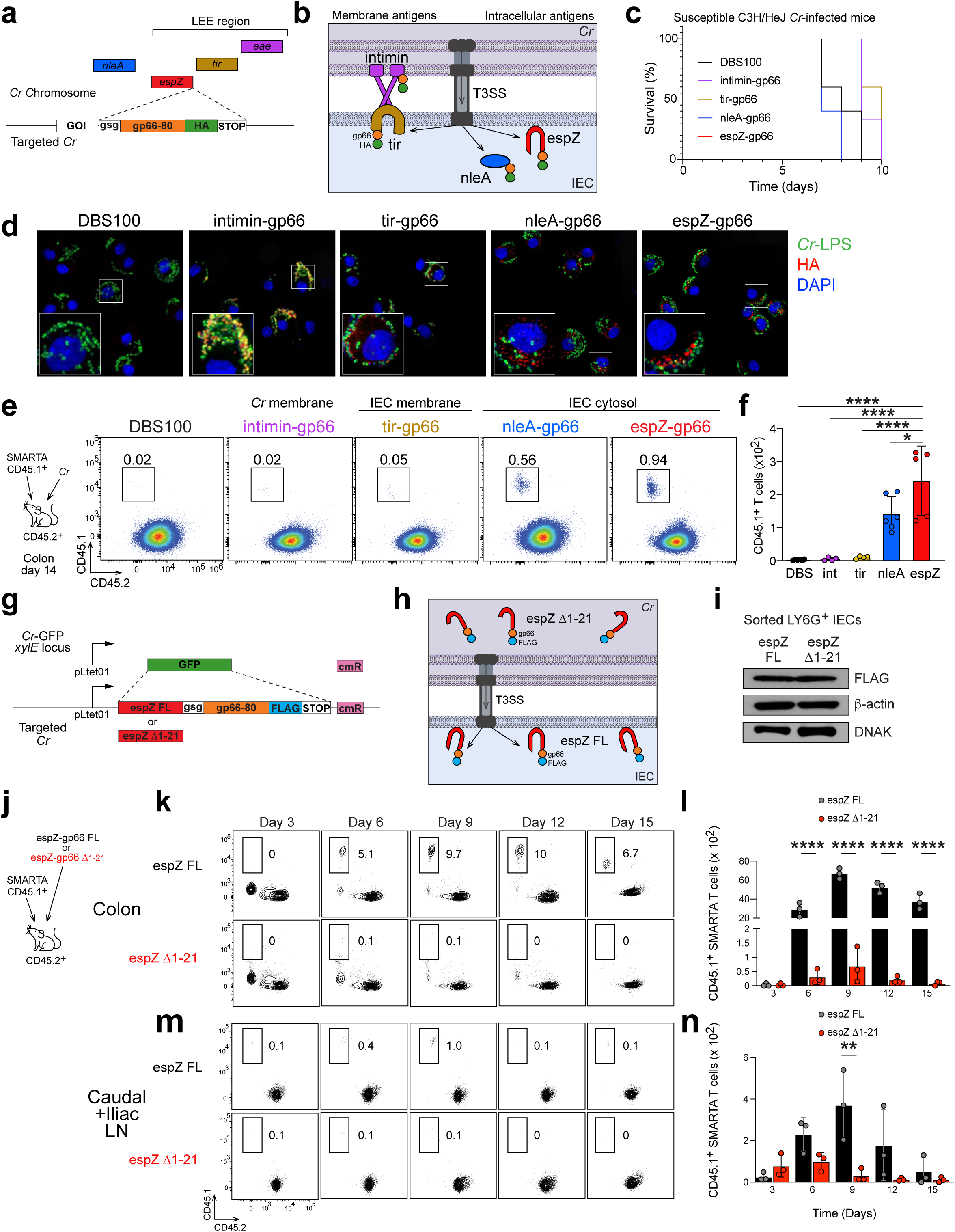
Subcellular localization of *Citrobacter rodentium* antigens determines magnitude of colonic CD4^+^ T cell responses. Candidate *Cr* proteins were tagged with LCMV gp66-80 epitope and used for adoptive transfer experiments with CD45.1^+^ SMARTA T cells. **a**, Schematic of gsg-gp66-80-HA construct inserted into *Cr* LEE-associated genes (*eae* (intimin), *tir*, and *espZ*) and non-LEE–associated gene *nleA*. **b**, Schematic of subcellular localizations of *Cr* effector proteins intimin, tir, nleA, and espZ relative to IEC-bound *Cr* cell. **c**, Survival of C3H/HeJ susceptible mice infected with DBS100 (WT), intimin-gp66, tir-gp66, nleA-gp66, and espZ-gp66. 4-5 mice per group; *n*=2 independent experiments. **d**, D8-infected EpCAM^+^GFP^+^ IECs were sorted from C57BL/6 mice infected with *Cr*-GFP (WT), intimin-gp66 *Cr*-GFP, tir-HA *Cr*-GFP, nleA-gp66 *Cr*-GFP, and espZ-gp66 *Cr*-GFP, were analyzed by immunofluorescence microscopy. **e-f**, Adoptively transferred naïve CD45.1^+^ SMARTA T cells into C57BL/6 mice infected with DBS100, intimin-gp66, tir-gp66, nleA-gp66, or espZ-gp66 at day 14 were stained for CD45.1 and CD45.2 and quantified by flow cytometry. 3-5 mice per group; *n*=2 independent experiments. One-way ANOVA; *p≤0.05 and ****p≤0.0001. **g**, Schematic of espZ FL-gp66-FLAG and espZ D1-21-gp66-FLAG constructs used to replace GFP coding sequence in *xylE* pseudogene in *Cr*-GFP parent strain. **h**, Schematic of subcellular locations of full-length espZ-gp66 translocated into the host IEC or espZ D1-21-gp66 protein, which remains associated with *Cr* bacteria. **i**, Total protein lysate from sorted mid-distal Ly6G^+^ DCCs from mice infected with espZ FL or espZ D1-21 were analyzed by Western blot for FLAG-tagged antigen with b-actin and DNAK as loading controls. **j**, Schematic of adoptively transferred naïve CD45.1^+^ SMARTA T cells into C57BL/6 mice infected with either espZ FL or espZ D1-21. **k-n**, On days 3-15 post-infection cell recovered from from colon (**k-l**) or caudal/iliac lymph nodes (**m-n**) were stained for CD45.1 and CD45.2 and analyzed by flow cytometry. 3-5 mice per group; *n*=2 independent experiments. One-way ANOVA; **p≤0.01, ***p≤0.001 and ****p≤0.0001.

Confocal immunofluorescence microscopy of infected colonocytes isolated at day 8 post-infection confirmed expected subcellular distributions of HA-tagged antigens (**Fig. 1d**)^24–29^. Intimin-gp66 was strictly associated with the bacterial cell surface, while tir-gp66 formed distinct puncta at sites of bacterial attachment to the apical plasma membrane of *Cr*-bound colonocytes. In contrast, both nleA-gp66 and espZ-gp66 were delivered into the host cell cytoplasm, with nleA showing more diffuse cytosolic distribution and espZ exhibiting intracytoplasmic clustering in a pattern suggestive of its partial association with bacterial pedestals^29^.

Using a modified adoptive transfer protocol to track responses of antigen-naive, congenically-marked SMARTA T cells, recipient mice were challenged with the different *Cr* strains (**Fig. 1e,f** and **ED Fig. 1d-i**). We observed striking differences in SMARTA T cell numbers recovered from the infected mid-distal colon at the peak of the mucosal T-cell response (day 14 post-infection; **Fig. 1e-f**). While both bacterial (intimin-gp66) and cEC (tir-gp66) membrane-localized antigen failed to induce accumulation of SMARTA cells that differed significantly from control wild-type (WT) *Cr*, both cytosol-delivered antigens (nleA and espZ) induced robust mucosal responses: nleA-gp66 and espZ-gp66 induced 28-fold and 47-fold greater SMARTA T cell accumulation in the colon, respectively, compared to intimin-gp66. Thus, differential compartmentalization of bacterial antigen had profound effects on the magnitude of the mucosal antigen-specific CD4 T cell response.

To determine if antigen abundance might be the basis for differential responses to the different *Cr* strains, we quantitated expression of the different HA-tagged proteins *in vitro* and *in vivo* (**ED Fig. 1j-l**). Western blot analysis of *in vitro* cultures of *Cr* activated to express virulence proteins revealed that intimin was detected in comparable amounts to tir and nleA in bacterial cell pellets, but was absent from culture supernatants, confirming its localization to the bacterial membrane. Tir and nleA appeared in both bacterial cell pellets and culture supernatants, reflecting their secretion by *Cr*, with tir being the most abundant secreted protein *in vitro*. EspZ protein was not detected *in vitro*, consistent with previous reports that this effector is poorly induced *in vitro*^30^.

Antigen expression *in vivo* was quantitated in *Cr*-DCC conjugates isolated from infected mice after engineering the four epitope-tagged variants into a GFP-expressing background strain of *Cr* (*Cr*-GFP) to enable flow cytometric sorting^1,31^. Intimin and espZ were detected at significantly lower levels than tir or nleA, which were the most abundant proteins in *Cr*-DCC conjugates of those assessed (**ED Fig. 1l**). Notably, both nleA and espZ induced similar Ag-specific responses despite the lower abundance of espZ. The observation that expression of Tir was greatest yet still induced a minimal mucosal SMARTA T cell response suggested that, while antigen abundance was important for immune recognition, antigen compartmentalization was a more critical determinant of immunogenicity. Together, these findings suggested that translocation of *Cr* bacterial antigens into the cytosol of infected cECs markedly enhanced the recruitment and/or retention of Ag-specific CD4 T cells in the *Cr*-infected colon.

To more directly determine whether translocation of bacterial antigens into the host cell cytosol drives optimal T cell responses, we examined the expression and SMARTA cell responses of full-length espZ and a truncated variant lacking the T3SS secretion signal. Minigenes that encoded expression of gp66 and a FLAG peptide fused to the carboxy terminus of either full-length espZ (espZ FL) or a truncated espZ variant lacking the N-terminal 21 amino acids (espZ Δ1-21) were generated and targeted to the *xylE* pseudogene locus of *Cr*^31,32^ (**Fig. 1g,h** and **ED Fig. 1a-c**), which drives high-level expression independently of the endogenous *espZ* locus, thereby avoiding perturbation of *Cr* pathogenicity (**ED Fig. 1a** and **data not shown**). Importantly, Western blotting of isolated *Cr*-bound cECs established that full-length espZ-gp66 and the Δ1-21 mutant proteins were comparably abundant in *Cr*-DCC conjugates, demonstrating that differences in epitope abundance should not confound the SMARTA T cell response (**Fig. 1i**).

Recipients of SMARTA T cell transfers challenged with the two espZ-gp66 *Cr* variants showed a marked bifurcation of the Ag-specific T cell response: The espZ FL strain induced a >100-fold increase in mucosal SMARTA T cells compared to the espZ Δ1-21 strain at the peak of the response (day 9; **Fig. 1j-l**). This did not appear to be due simply to different retention of the Ag-specific cells in the infected mucosa, as kinetic analysis of SMARTA cell numbers in the lymph nodes (LNs) draining the distal colon (caudal and iliac LNs) showed comparable clonal expansion differences throughout the infection time course (**Fig. 1m,n**). Collectively, these results indicate that T3SS-mediated delivery of antigens into the IEC cytosol induces CD4 T cell responses of much greater amplitude than those of bacterially- or IEC cell membrane-associated antigens, even when controlled for the same T cell epitope and quantity of available antigen.

### T cell priming is initiated in cecum-draining lymphoid tissue before spread to distal colon

*Cr* cultured *in vitro* is known to colonize the cecum—where expression of pathogenicity genes required to assemble the T3SS begins^17,33,34^—prior to spread downstream to distal colon^32,88^, yet the precise localization and immunological consequences of this early colonization are incompletely understood. Absent models to track clonal T cell responses to *Cr* antigens, it has not been possible to define sites where these responses are initiated. In view of the foregoing data indicating a dominant role for injected cytosolic antigen in the distal colon and the LNs draining it, we sought to determine where Ag-specific T-cell responses were initiated and whether the relatively muted response to non-cytosolic antigens might reflect altered geography of T-cell priming—in the cecum rather than distal colon.

Using fluorescence in situ hybridization (FISH) with probes for total Eubacteria and *Cr*, respectively, we examined bacterial localization during cecal colonization early in infection in tissue processed to preserve mucus architecture (**Fig. 2a-c**). Remarkably, *Cr* exhibited highly focal epithelial colonization in the region of the cecal apex, in close proximity to—but not in direct association with—the cecal patch, a GALT structure analogous to Peyer’s patches in the small intestine^35^ (**Fig. 2c-f**). *Cr* was also abundant within the cecal mucus layer, as well as free in the lumen. The basis for selective colonization of this restricted region of cecal epithelium is unknown but its proximity to the cecal patch raised the possibility that this mucosal lymphoid structure might serve as an early site of antigen sampling and T cell priming^36^. We therefore tracked SMARTA T cell dynamics following infection with the *Cr* strains espZ-gp66 or intimin-gp66 across regions colonized by *Cr*—cecum and its draining distal mesenteric lymph node (dMLN) as well as the distal colon and its draining LNs (caudal and iliac LNs) (**Fig. 2g-r**).

**Fig. 2:**
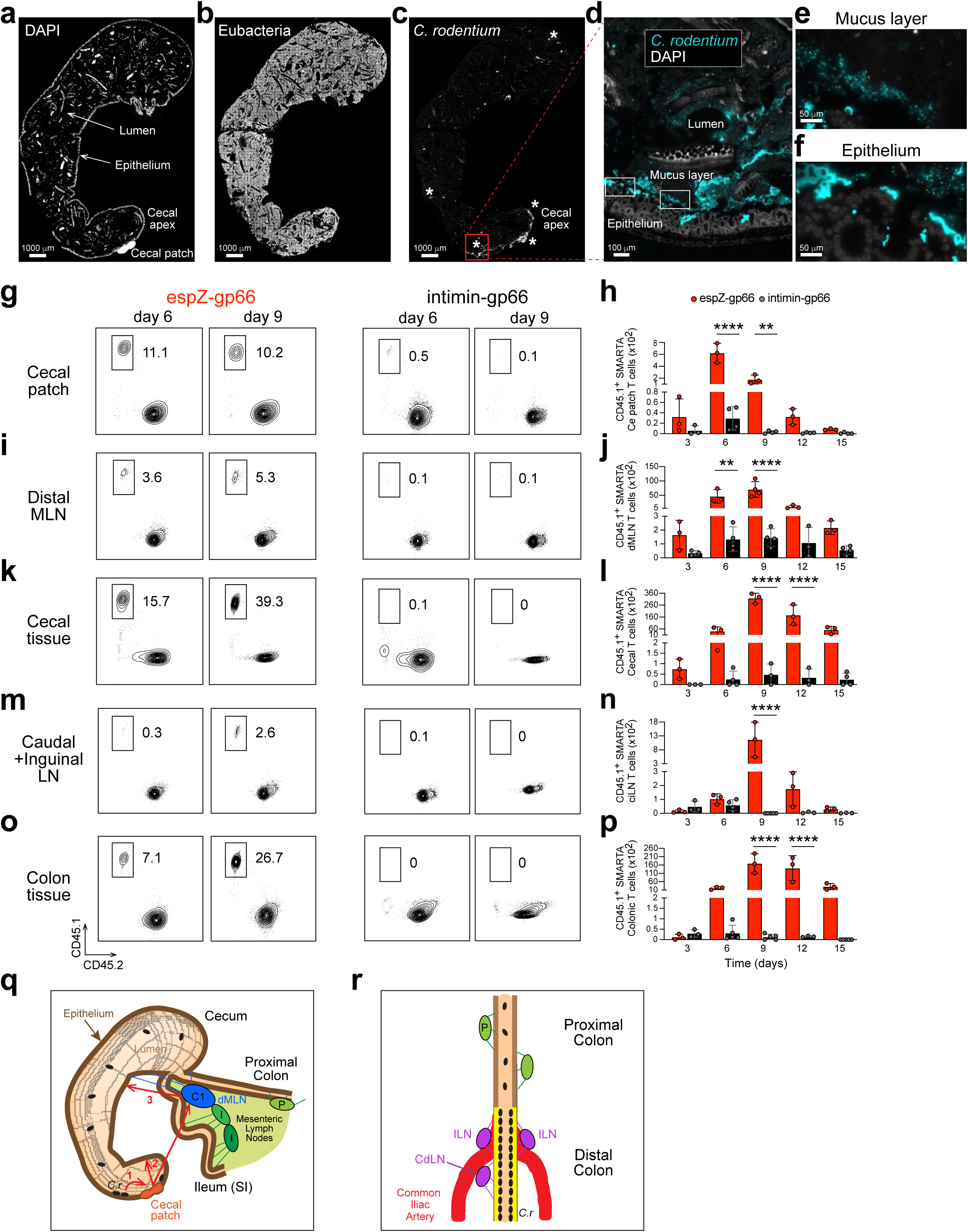
Early *C. rodentium* colonization in cecum initiates CD4^+^ T cell priming in gut-associated lymphoid tissue. Ceca were collected from 48-hour DBS100-infected C57BL/6 mice and fixed in ice-cold methacarn solution to preserve mucus layer structure. Cecal sections were stained with DAPI (**a**) and FISH probes for total Eubacteria (**b**) and *Cr* (**c-f**). Adoptively transferred naïve CD45.1^+^ SMARTA T cells in C57BL/6 mice infected with espZ-gp66 or intimin-gp66 were analyzed by flow cytometry on days 6 and 9 post-infection from cecal patch (GALT, **g-h**), distal mesenteric lymph node (dMLN, **i-j**), cecal tissue (**k-l**), caudal and iliac lymph nodes (ciLN, **m-n**), and colon tissue (**o-p**). 3-5 mice per group; *n*=2 independent experiments. One-way ANOVA; *p≤0.05, **p≤0.01, ***p≤0.001 and ****p≤0.0001. **q,r**, Schematics depicting initial antigen acquisition via cecal patch and dMLN prior to cecal tissue infiltration (**q**) or colonic attachment of *Cr* to distal colon and antigen drainage into ciLN following initial cecal colonization (**r**).

In response to espZ-gp66, SMARTA cell numbers were greatest in the cecal patch day 6 post-infection before rapidly declining by day 9 (**Fig. 2g-h**). In the cecum-draining dMLN, the SMARTA response showed similar kinetics, with substantial SMARTA cell accumulation beginning at day 6, peaking at day 9, and rapidly declining by day 12 (**Fig. 2i-j**). In contrast, significant T cell infiltration into the cecal mucosa was delayed until day 9 and remained elevated through day 12 before declining (**Fig. 2k-l**). Notably, although the response in cecal patch was modestly accelerated compared to dMLN, the peak amplitude was considerably greater (∼10-fold) in dMLN and the number of Ag-specific cells recovered from the cecal mucosa was greater still (∼5-fold that of dMLN), suggesting that priming of the response in cecal patch—presumably generated in response to bacterial antigens translocated directly across the patch epithelium since there was no detectable colonization of patch itself—is minor compared to that generated in dMLN. The clonal response in the colon mucosa (**Fig. 2m-p**) largely tracked with that observed in cecum, whereas the detectable response in the dMLN preceded that of distal colon-draining LNs and was of considerably greater amplitude (> 5-fold) at the peak, indicating that the primary site for induction of the T cell response is the lymph node that drains the cecum—consistent with the initial expression of translocated virulence proteins by *Cr* in the cecum.

The response to intimin-gp66 was considerably weaker across all tissue sites and time points and, although measurable in draining LNs, was essentially undetectable in the cecal and distal colonic mucosae (**Fig. 2g-p** and **Fig. 1e,f**), indicating that the differences in responses to espZ-gp66 and intimin-gp66 could not be ascribed to differing sites of induction. Accordingly, in a survey of the dynamics of SMARTA cells induced by infection with either espZ FL or Δ1-21 espZ that was extended to include the cecum and all draining LNs (**ED Fig. 2**), the magnitude of the clonal response to injected antigen dwarfed that of the same antigen sequestered in the bacterial cytosol—at each time point and tissue site—mirroring our findings comparing espZ-gp66 and intimin-gp66. Notably, the kinetics of the T-cell response to espZ FL was not appreciably accelerated in the cecum despite its constitutive expression of espZ from the *xylE* gene locus, consistent with rapid expression of native espZ on arrival of Cr in the cecum. Also, the greatest numbers of SMARTA cells were recovered from the infected mucosae, consistent with their retention there by antigen recognition on the infected epithelium. Collectively, these results established that significantly weaker responses to bacterially-associated antigen—both surface and cytosolically localized—were consistent across the cecal and distal colonic sites of *Cr* colonization, and not simply due to regional differences in antigen delivery dynamics. Rather, the subcellular localization of antigen was the major determinant of the magnitude of the T-cell response, which appeared to be initiated in the cecum-draining lymph node where injected antigens were first available from a site in the cecal apex before tracking with the spread of antigen to the epithelium of the distal colon and the lymph nodes that drain it. Thus, the quality of antigen superseded quantity of antigen as the major determinant of the magnitude and distribution of clonal T cells responding to *Cr* antigens.

### Epithelial cell presentation of injected bacterial antigens promotes CD4 Trm cell development

Despite considerable speculation, definitive evidence supporting an important role for non-professional APCs in the development of memory T cells has been limited^37–40^. In view of the superior ability of injected antigens to drive mucosal CD4 T cell responses during acute infection and our previous finding of antigen presentation by IECs as a key mechanism for recruiting CD4 T cell help^1^, we speculated that T cell recognition of injected bacterial antigens presented by IECs might directly contribute to the development of mucosal memory T cells. Assessment of total CD69^+^CD103^−^ and CD69^+^CD103^+^ SMARTA Trm cell numbers generated in response to different *Cr* strains revealed striking differences in both number and phenotype (**Fig. 3a-e**). At day 42 post-infection, more than 3 weeks after bacterial clearance, few SMARTA Trm cells were generated in response to intimin-gp66. Notably, despite the modest mucosal T cell response induced by tir-gp66 (**Fig. 1a-f**), there was significant induction of both CD69^+^CD103^−^ and CD69^+^CD103^+^ Trm cells. However, nleA-gp66 and espZ-gp66 induced far greater numbers of SMARTA Trm cells (22- and 30-fold higher, respectively, than intimin-gp66), which represented the majority of the total recovered SMARTA cells (**Fig. 3d-e**). Thus, Trm cell development was strongly biased towards injected bacterial antigens, especially those localized to the IEC cytosol, but also including a weaker response to the membrane-embedded variant, tir. Notably, the ratios of CD103^−^ to CD103^+^ fractions were similar across these antigenic variants, suggesting that the relative induction of these subsets was independent of epithelial antigen compartmentalization.

**Fig. 3:**
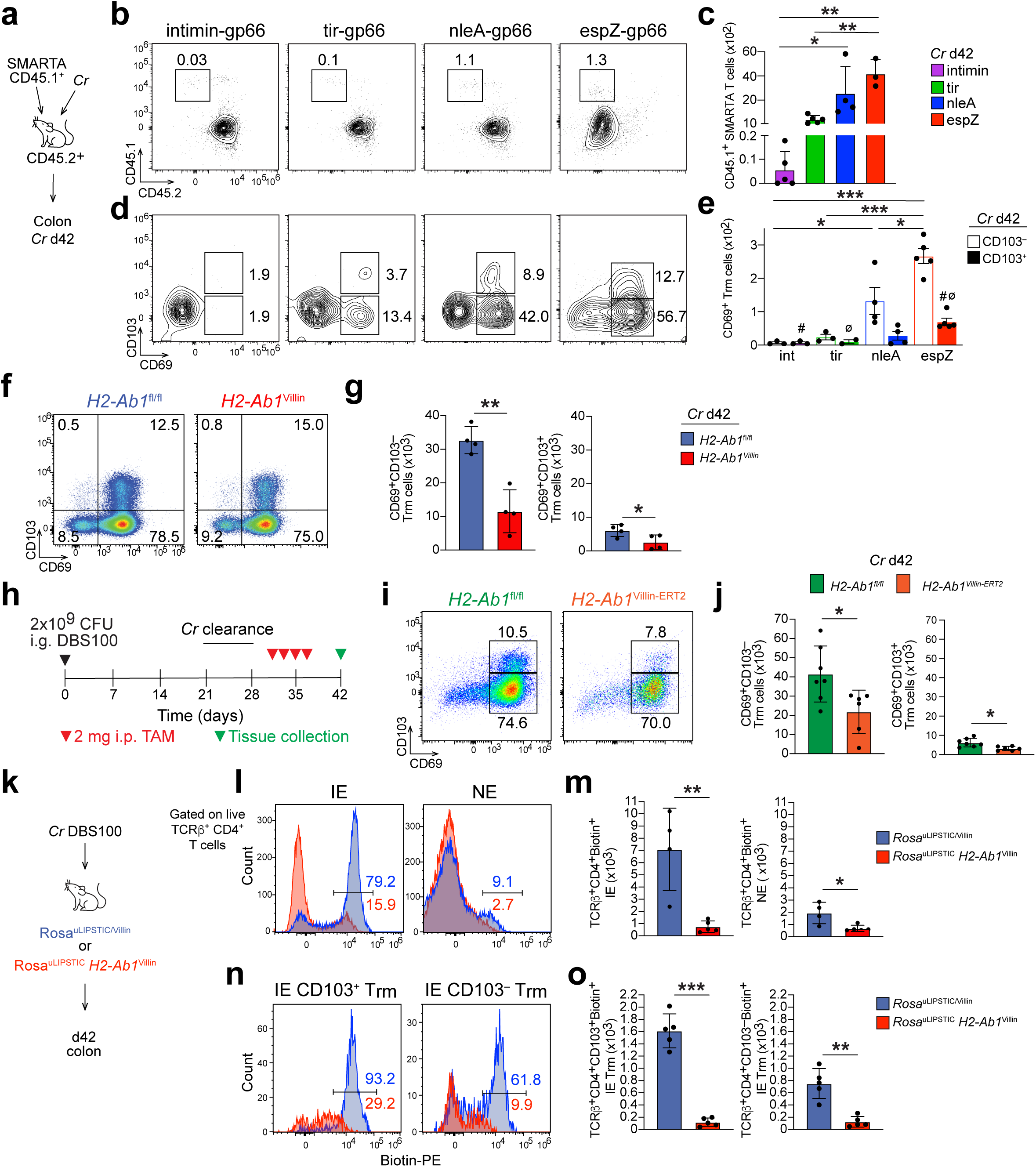
Epithelial cell presentation of injected *C. rodentium* antigens drives CD4^+^ tissue-resident memory T cell formation and maintenance. Naïve CD45.1^+^ SMARTA T cells were adoptively transferred into C57BL/6 mice infected with either intimin-gp66, tir-gp66, nleA-gp66, or espZ-gp66 *Cr* strains (**a**). **b,c**, Cells from colons of mice were collected on day 42 post-infection, stained for CD45.1 and CD45.2 and analyzed by flow cytometry. 3-5 mice per group; *n*=2 independent experiments. One-way ANOVA; *p≤0.05 and **p≤0.01. **d,e**, Colonic cells from d42-infected mice were stained for CD4, TCRβ, CD69, CD103, and analyzed by flow cytometry to quantitate numbers of CD69^+^CD103^−^ and CD69^+^CD103^+^ Trm cells. One-way ANOVA; *p≤0.05 and ***p≤0.001 (CD103^−^ with different antigens) and ^#^p≤0.05 and ^Ø^ p≤0.05 (CD103^+^ with different antigens). **f,g**, Cells from colons of DBS100-infected *H2-Ab1*^fl/fl^ (blue) and *H2-Ab1*^Villin^ (red) mice were collected on day 42 stained as in **d** and analyzed by flow cytometry to quantitate the number of CD69^+^CD103^−^ and CD69^+^CD103^+^ Trm cells. 3-5 mice per group; *n*=2 independent experiments. Two-tailed unpaired *t*-test; *p≤0.05 and **p≤0.01. **h**, Schematic of tamoxifen (TAM)-induced deletion of *H2-Ab1* post-*Cr* clearance in *H2-Ab1*^fl/fl^ and *H2-Ab1*^Villin-ERT2^ mice receiving TAM days 33-36 post-infection. **i-j**, Cells from colons of DBS100-infected *H2-Ab1*^fl/fl^ (green) and *H2-Ab1*^Villin-ERT2^ (orange) mice were stained as in **d** and analyzed by flow cytometry on d42 post-infection to quantitate number of CD69^+^CD103^−^ and CD69^+^CD103^+^ Trm cells after TAM-induced deletion of *H2-Ab1*. 5-6 mice per group; *n*=2 independent experiments. Two-tailed unpaired *t*-test; *p≤0.05 and **p≤0.01. **k**, Schematic of ‘Labelling Immune Partnerships by SorTagging Intercellular Contacts’ (LIPSTIC) experiments using Rosa^uLIPSTIC/Villin^ (blue) and Rosa^uLIPSTIC^*H2-Ab1*^Villin^ (red) mice at day 42 post-infection following biotin-LPETG administration. **l,m**, Cells isolated from intraepithelial (IE) or non-epithelial (NE) compartments of colon from Rosa^uLIPSTIC/Villin^ (blue) and Rosa^uLIPSTIC^*H2-Ab1*^Villin^ (red) mice at day 42 post-infection were stained as in **d** with SA and numbers of biotin^+^ CD4^+^ cells were quantified in IE and NE compartments (**l,m**) as well as the number of biotin+ CD69^+^CD103^−^ and CD69^+^CD103^+^ Trm cells in the IE compartment (**n,o**). 3-5 mice per group; *n*=2 independent experiments. Two-tailed unpaired *t*-test; *p≤0.05 and **p≤0.01.

Preferential generation of Trm cells by injected bacterial antigens suggested that epithelial cell antigen presentation might be critical for Trm cell programming. We found previously that epithelial MHCII is upregulated during acute *Cr* infection, is critical for controlling bacterial burden and pathology, and that loss of epithelial MHCII leads to decreased numbers of antigen-specific effector CD4 T cells^1^. To determine whether epithelial MHCII is required for Trm induction, we compared control *H2-Ab1*^fl/fl^ mice with *H2-Ab1*^Villin^ mice that have deficiency of MHCII specifically targeted to IECs. Compared to control *H2-Ab1*^fl/fl^ mice, *H2-Ab1*^Villin^ mice showed significantly reduced CD69^+^CD103^−^ and CD69^+^CD103^+^ CD4 Trm populations at day 42 post-infection (**Fig. 3i-j**), demonstrating that epithelial MHCII is necessary for optimal CD4 Trm development. To determine whether ongoing epithelial MHCII expression is required to maintain Trm cells after bacterial clearance, we employed tamoxifen-inducible deletion of epithelial MHCII (*H2-Ab1*^Villin-ERT2^ mice) (**Fig. 3k**). Loss of MHCII expression by IECs in the memory phase resulted in significant reduction of both CD69^+^CD103^−^ and CD69^+^CD103^+^ Trm subsets (**Fig. 3l,m**), indicating that continued epithelial MHCII expression contributed to Trm maintenance after pathogen clearance. Thus, presentation of injected bacterial antigen to CD4 T cells is required for optimal induction of CD4 Trm cell development and sustained MHCII expression by IECs is important for maintenance of these cells.

To extend these findings, we assessed MHCII-dependent interactions between cECs and CD4 T cells during the memory phase following resolution of *Cr* infection, employing the LIPSTIC (Labeling Immune Partnerships by SorTagging Intercellular Contacts) transgenic system^41,42^, which mediates unidirectional labeling of lineage-specific Cre-activated donor cells to any recipient cell in close proximity (**Fig. 3n**). In *Rosa*^uLIPSTIC/Villin^ mice, where epithelial cells can transfer biotin to interacting partners, 79% of recovered intraepithelial CD4 Trm cells were biotin-positive at day 42 post-infection (**Fig. 3o,p**). In contrast, *Rosa*^uLIPSTIC^/*H2-Ab1*^Villin^ mice, which are devoid of MHCII-dependent interactions, showed markedly reduced biotin labeling (16% of recovered IE CD4 Trm cells recovered), reflecting the ∼5-fold reduction in total biotin-labelled intraepithelial CD4 T cells. There was modest, MHCII-dependent labelling of non-epithelial CD4 T cells, indicating detectable but relatively infrequent interactions between these T cells and IECs than those observed for intraepithelial CD4 T cells—which were largely lost in absence of IEC MHCII. Notably, although colonocyte–Trm cell interactions were less frequent for CD103^−^ CD4 Trm cells—despite their greater abundance within the colonic epithelium—they were nevertheless substantial (∼62% biotin+), indicating that the majority of both CD103^−^ and CD103^+^ Trm subsets sustain ongoing interactions with cECs that are critically dependent on MHCII-dependent interactions. Thus, MHCII-dependent interactions required for the induction and/or maintenance of *Cr*-induced CD4 Trm cells reflect direct IEC-Trm cell physical interactions and indicate that sustained contacts between cECs and both CD4 Trm cell subsets. Collectively, these results establish a dominant role for colonocyte presentation of injected bacterial antigens in promoting the development of both major subsets of mucosal CD4 Trm cells. They support a model in which ongoing MHCII-dependent direct interactions between IECs and Trm cells are required to sustain the Trm population. Moreover, these findings establish that sustained direct physical interactions between cECs and Trm cells within the colonic epithelium are favored for CD103^+^ Trm cells, but are substantial for both CD103^−^ and CD103^+^ subsets.

### Antigen localization dictates spatial organization of memory CD4 T cells

The foregoing results established a marked quantitative bias in favor of effector and memory CD4 T cells responding to bacterial antigens injected into the cytosol of IECs during *Cr* infection but did not define the microanatomical disposition of these cells. We therefore employed light sheet fluorescence microscopy (LSFM) to visualize and quantitate the spatial distribution of Ag-specific memory T cells contingent on the localization of recognized antigen. Following infection of SMARTA T cell transfer recipients with intimin-gp66 or espZ-gp66 *Cr*, striking differences in the localization of Ag-specific CD4 T cells were evident contingent on response to bacterial- versus cEC-associated antigen.

In response to intimin-gp66 *Cr*, SMARTA T cells were extremely rare in the epithelium, with only approximately 10 CD45.1^+^ cells detected throughout the imaged tissue volume (**Fig. 4a**). Remarkably, no SMARTA cells were identified in the colonic lamina propria (LP). Instead, most SMARTA T cells (93% of a total of approximately 137 cells) localized to organized gut-associated lymphoid tissue (GALT; colonic patches and isolated lymphoid follicles, or ILFs). Thus, the low-amplitude clonal response to bacterially associated antigen strongly favored localization to intramucosal lymphoid tissues (GALT) rather than the epithelial barrier during the memory phase.

**Fig. 4:**
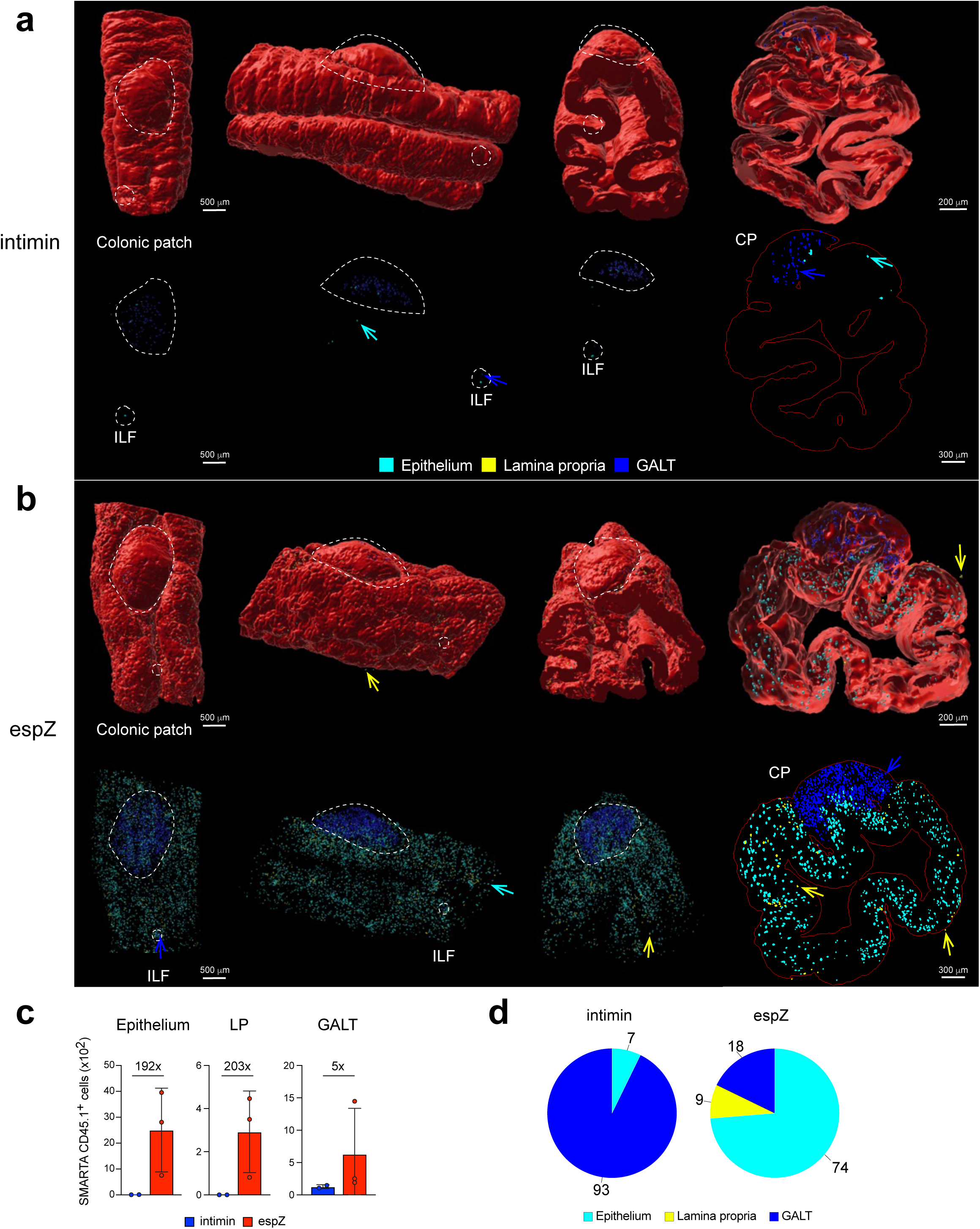
Differential antigen localization promotes distinct patterns of CD4 T cell memory. Light-sheet fluorescence microscopy (LSFM) was performed on day 42 colons from mice that received naïve CD45.1^+^ SMARTA T cells and infected with either intimin-gp66 or espZ-gp66 strains. Mice received i.v. anti-CD45.1 and anti-EpCAM antibodies prior to tissue collection and tissue clearing. Machine learning was used to assign CD45.1^+^ cells into epithelium (≤ 0 mm from EpCAM^+^ cells; cyan), lamina propria (LP; > 0 mm from EpCAM^+^ cells; yellow), or GALT (manually assigned based on tissue architecture; blue). **a,b**, Images of CD45.1^+^ cells in colonic patch and isolated lymphoid follicles (demarcated by dashed white line) from mid-distal colon of intimin-gp66 infected (**a**) or espZ-gp66 infected (**b**) C57BL/6 mice. **c**, SMARTA cell numbers in different anatomical locations calculated from LSFM images. **d**, Pie charts of percentages of CD45.1^+^ SMARTA cells within indicated tissue compartments of mice infected with intimin-gp66 or espZ-gp66.

In stark contrast, infection with espZ-gp66 *Cr* generated a far greater population of intraepithelial memory SMARTA cells, with ∼2,500 Trm cells distributed within the epithelial compartment—a 250-fold increase compared to intimin-specific cells (**Fig. 4b**). The GALT contained relatively fewer SMARTA cells (629)—representing a reciprocal distribution compared with that induced by intimin-gp66. Moreover, the number of epithelium-associated Trm cells significantly outnumbered those found in the lamina propria (∼8-fold increase). Notably, relatively few SMARTA memory T cells were retained in the mucosal lamina propria (9% of the total, or 300 cells), despite the preponderance of cells in this niche at the height of the effector response to infection (**see below** and **data not shown**), suggesting that signals required to sustain T cells in this anatomic niche are limited. These imaging data provide compelling visual evidence that antigen compartmentalization determines not only the magnitude but also the microanatomical positioning of memory CD4 T cells. The dense network of memory T cells distributed throughout the epithelial layer emphasize that injected antigens preferentially establish frontline memory defense at the barrier surface—poised at the site they are likely to re-encounter the same antigen in subsequent infections.

### Epithelial cell antigen presentation underpins bidirectional epithelial-T cell crosstalk

The discovery of a specialized role for IEC presentation of injected antigens in driving the magnitude, developmental trajectory and tissue distribution of the CD4 T-cell response led us to examine the bidirectional signals exchanged between the antigen-presenting DCCs and responding T cells. To comprehensively characterize epithelial-T cell information exchange mediated by IEC antigen presentation, we performed single-cell RNA sequence (scRNA-seq) analysis on sorted EpCAM^+^ epithelial cells and CD45^+^ immune cells from the intraepithelial (IE) and non-epithelial (NE) compartments of the mucosa of naïve and infected *H2-Ab1*^fl/fl^ and *H2-Ab1*^Villin^ mice (**ED Fig. 4a,b**). This has provided an unprecedented overview of the transcriptional profiles of epithelial and immune cell subpopulations during and following infection by *Cr* (**ED Fig. 4a,b and Supplementary Table 2**). The effects of targeted deficiency of MHCII expression by IECs and consequent loss of antigen-dependent dialog with CD4 T cells were substantial. Although primarily impacting the direct reciprocal interactions between IECs and *Cr*-specific CD4 T cells that are lost due to failed antigen presentation, the impact extended to other epithelial and immune cell populations—both innate and adaptive.

Analysis of *H2-Ab1* (MHCII) expression across all clusters revealed that while professional mucosal antigen-presenting cells (e.g., DCs and macrophages) maintained high expression of MHCII at all timepoints, specific epithelial cell subpopulations upregulated MHCII in response to infection (**Fig. 5a and ED Fig. 5**). Temporal analysis showed that the greatest expression of *H2-Ab1* by epithelial cells was found at day 14 of infection in distal colonocytes (DCCs)—particularly pro-DCCs, mature DCCs, and pro-senescent CCs (**Fig. 5b**). Notably, MHCII expression persisted in multiple colonocyte subsets at day 42, including pathogen-induced colonocytes (P-I CCs), pro-DCCs, mature DCCs and pro-senescent CCs—well beyond the clearance of *Cr* and resolution of effector T cell infiltrates and innate immune cell infiltrates in the infected colonic lamina propria (**Fig. 4** and **data not shown**). The persistent surface expression of MHCII was confirmed by flow cytometric and independent transcriptomic analysis (**Fig. 5c-e**).

**Fig. 5:**
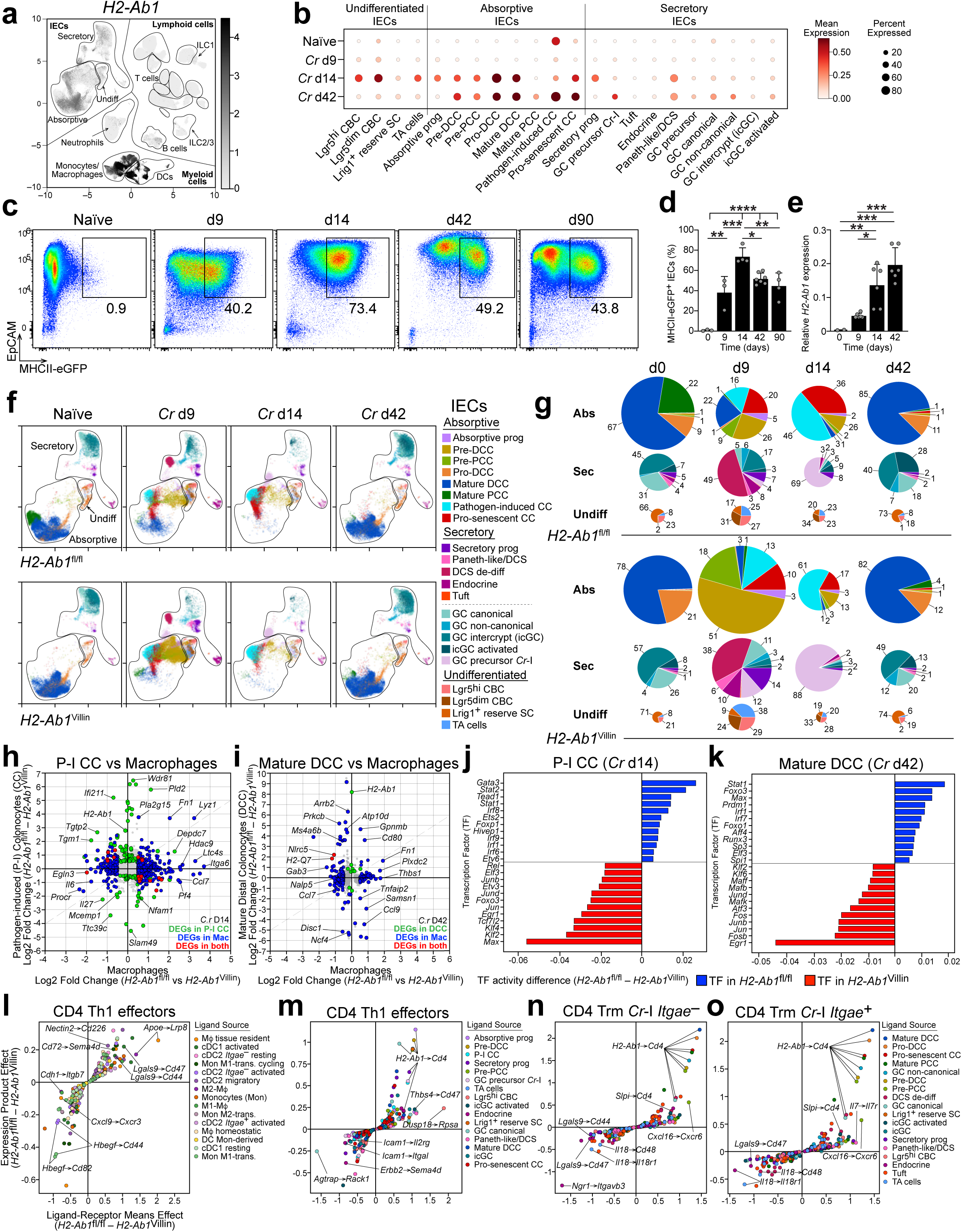
Epithelial MHCII expression sustains IEC-T cell communication during and following *C. rodentium* infection. scRNA-seq was performed on EpCAM^+^ and CD45^+^ cells isolated from intraepithelial (IE) and non-epithelial (NE) compartments from mid-distal colons of *H2-Ab1*^fl/fl^ and *H2-Ab1*^Villin^ mice that were uninfected (naïve) or *Cr*-infected for 9, 14 or 42 days (n=2). **a**, UMAP analysis of *H2-Ab1* expression in key labeled clusters (see **ED Fig. 4** for detailed demarcation of clusters). **b**, Dot plot of *H2-Ab1* expression in epithelial cell clusters in naïve and day 9, 14 or 42 *Cr*-infected mice. **c,d**, EpCAM^+^ IECs isolated from naïve or day 9, 14, 42 or 90 *Cr*-infected MHCII-eGFP mice were analyzed by flow cytometry. 3-5 mice per group; *n*=2 independent experiments. One-way ANOVA; *p≤0.05, **p≤0.01, ***p≤0.001 and ****p≤0.0001. **e**, *H2-Ab1* gene expression in EpCAM^+^ IECs sorted from naïve or day 9, 14 or 42 of *Cr*-infected C57BL/6 mice determined by RT-PCR. 3-5 mice per group; *n*=2 independent experiments. One-way ANOVA; *p≤0.05, **p≤0.01, and ***p≤0.001. **f**, UMAPs of IEC clusters from naïve and day 9, 14 or 42 *Cr*-infected *H2-Ab1*^fl/fl^ and *H2-Ab1*^Villin^ mice. **g**, Pie charts of the percentages of absorptive, secretory, and undifferentiated IEC clusters from the indicated time points and mouse genotypes. Pie chart areas normalized to largest pool of cells (d9 infected *H2-Ab1*^Villin^ mice). **h,i**, Two-way scatter plot of DEGs from P-I CCs vs DEGs from pooled *H2-Ab1*^+^ macrophage clusters from day 14 (**h**) and DEGs from mature DCCs vs DEGs from pooled *H2-Ab1*^+^ macrophage clusters from day 42 (**i**) scRNA-seq data. **j,k** pyScenic TF regulon analysis performed in P-I CCs at day 14 (**j**) and Mature DCCs at day 42 (**k**). **m-o**, Ligand-receptor analysis performed via LIANA on myeloid-CD4 Th1 effector interactions (d14, **l**), IEC-CD4 Th1 effector interactions (d14, **m**), IEC-CD4 CD103^−^ Trm interactions (d42, **n**), and IEC-CD4 Trm CD103^+^ interactions (d42, **o**).

Comparison of epithelial cell developmental trajectories between *H2-Ab1*^fl/fl^ and *H2-Ab1*^Villin^ mice highlighted altered colonocyte dynamics during active infection in absence of epithelial MHCII (**Fig. 5f,g**). With the exception of a relative increase in fraction of proximal colonocytes (PCCs) in *H2-Ab1*^fl/fl^ mice that appears to reflect sampling bias (**data not shown**), no significant differences in CC populations were observed between the genotypes at homeostasis, reflecting a lack of colonocyte expression of MHCII at steady-state. However, *H2-Ab1*^Villin^ mice failed to show the normal increase in shedding of pro-senescent colonocytes during active infection, consistent with our previous finding of a critical roles for T cell-derived IL-22 in programming these cells for removal to counter *Cr* effector proteins that promote DCC retention^1^ (**Fig. 5g**). This was evident by a decrease in both the relative and absolute number of pro-senescent CCs and an increase in total number of DCCs in *H2-Ab1*^Villin^ mice at the peak of bacterial load in WT mice (day 9). Notably, the total number of DCCs in *H2-Ab1*^Villin^ mice contracted at day 14, consistent with impaired recruitment of T cell-derived IFNγ signaling, which is a key driver of increased absorptive IEC proliferation during infection^44^ (see below).

Although there were shifts in the relative proportions of total absorptive versus secretory lineage cells consequent to changes in the pool size of MHCII-deficient absorptive IECs, the composition of secretory lineage subsets was less altered, indicating that, beyond the global shift in relative and absolute numbers of secretory and absorptive lineage cells in favor of the latter, intralineage shifts in secretory cells were not as impacted as those in the absorptive lineage—likely reflecting the absence of antigen loading of secretory cells by injected *Cr* proteins, although this will require further study. Nevertheless, an increase in GC precursor *Cr*-I cells was observed in *H2-Ab1*^Villin^ mice, which may reflect reduced T cell-derived IFNγ in absence of epithelial antigen presentation, as IFNγ normally drives contraction of the GC lineage in favor of the absorptive lineage.

Two-way scatter plots comparing genes that were differentially expressed between epithelial and myeloid populations confirmed that *H2-Ab1* deletion was specific to epithelial cells, with no significant effect on *H2-Ab1* expression by myeloid cells (**Fig. 5h-i**). Thus, while there were notable gene expression differences and significant shifts in the total number and subset composition of macrophages during active infection that were contingent on MHCII-dependent interactions between *Cr*-infected DCCs and CD4 T cells (e.g., loss of regulatory macrophages at day 9; **ED Fig. 6 and Supplementary Table 2**), differences in expression of *H2-Ab1* by *H2-Ab1*^fl/fl^ and *H2-Ab1*^Villin^ macrophage or dendritic cell populations was not significantly altered (**ED Fig. 5**), indicating that these mucosal professional APCs likely retained antigen-presenting functions. This was confirmed by gene expression patterns indicating that macrophage and DC subsets from both *H2-Ab1*^fl/fl^ and *H2-Ab1*^Villin^ mice showed comparable up-regulation of genes that are downstream of CD40 signaling (e.g., **Fig. 5h,i** and **data not shown**), and therefore contingent on direct T cell interactions during antigen presentation.

Transcription factor regulon analysis using pySCENIC^45^ revealed that IFNγ-responsive transcription factors, including Stat1, Stat2, Irf1, Irf7, and Irf8, were the most significantly enriched in infected *H2-Ab1*^fl/fl^ IECs (e.g., P-I CCs at day 14 and mature DCCs at day 42) compared to MHCII-deficient IECs (**Fig. 5j-k**). This implicated a major role for epithelial cell antigen presentation to the numerically dominant Th1 cell population in programming the DCC functional response both during and following active engagement with *Cr*. While the MHCII-dependent signals that maintain this program following bacterial clearance—and presumably clearance of bacterial antigens—are unclear, the sustained IFNγ signatures suggest that ongoing epithelial-T cell crosstalk is critical in dampening innate signaling programs in IECs that fail to extinguish in absence of T cell-dependent “instructions.” Most notably, induction of a Prmd1 regulon downstream of IFN signaling in IECs late in the response (day 14) is thought to be important in feedback restraint of pro-inflammatory IFNγ-driven signals that impede restoration of homeostatic IECs functions^46–48^. Prdm1 (Blimp-1) actions also promote a transcriptional environment that allows up-regulation of Foxo1 and Foxo3 and their re-entry into the nucleus^49^, where they can cooperate with Prdm1 in restraining actions downstream of innate sensing of *Cr* by NOD family intracellular signaling mediated by AP-1 and NF-κB family members^50^. Thus, major transcriptional networks activated or repressed contingent on recruitment of T-cell help by antigen-presenting DCCs appeared to be dominated by IFNγ-mediated signaling into IECs by *Cr*-specific Th1 cells. In absence of this antigen-dependent signaling from T cells, the innate response to inflammasome signals—evidenced by the observed enrichment of AP-1 family regulon signatures in colonocytes of *H2-Ab1*^Villin^ mice (e.g., Fos, Jun, Junb, Jund)^51–53^—failed feedback restraint that restores homeostasis. Notably, included among these signatures are members of the Klf family of transcription factors (e.g., Klf2, Klf4), which, like Prdm1 and Foxo factors act to repress AP-1 family activity in IECs^54^. Enrichments of Klf family regulons in *H2-Ab1*^Villin^ mice suggests that this feedback mechanism can be elicited in absence of T cell cytokine inputs.

In extension of these findings, we performed an agnostic ranking of ligand-receptor interactions between different between epithelial-T cell pairs using LIANA+ analysis^55^ to identify specific intercellular communication pathways. In agreement with our findings above (**Fig. 5h,i**), we found no differential H2-Ab1-Cd4 interactions between different myeloid cells and Th1 effector cells in *H2-Ab1*^fl/fl^ or *H2-Ab1*^Villin^ mice at day 14 of infection (**Fig. 5l**). This was in stark contrast to epithelial cell-Th1 cell interactions, where H2-Ab1–Cd4 interactions dominated the most differentially scored ligand-receptor interactions between *H2-Ab1*^fl/fl^ and *H2-Ab1*^Villin^ mice (**Fig. 5m**). This epithelial-specific MHCII engagement persisted at day 42, with strong *H2-Ab1* interactions between epithelial cells and both CD103^−^ and CD103^+^ CD4 Trm subsets observed only in *H2-Ab1*^fl/fl^ mice (**Fig. 5n-o**). Notably, additional enriched ligand-receptor pairs identified in *H2-Ab1*^fl/fl^ mice included Cxcl16-Cxcr6, which is important in supporting tissue residency of T cells in the colonic epithelium^56,57^ and Il-7–Il-7r, which supports T cell survival^58,59^. In this regard, we found significant shifts in the production of *Il7* and *Il15* transcripts by distinct subsets of IECs over the course of *Cr* infection that may impact the source of survival signals used by CD4 T cells as they transition to the memory compartment (**Fig. ED 7**). It is also notable that in absence of T-cell help to epithelial cells in *H2-Ab1*^Villin^ mice, we find Il-18-Cd48 and Il-18–Ir-18r1 signatures between IECs and T cells late after bacterial clearance in the memory phase. Ongoing *Il18* transcription would appear to reflect loss of resolution signals downstream of T cell cytokine signaling to IECs, resulting in sustained, unchecked innate danger signaling pathways activated by inflammasome–AP-1 programs that drive persistent epithelial IL-18 production, consistent with the regulon data above (**Fig. 5j,k**).

Reciprocal analysis of the scRNA-seq dataset focusing on how deficiency of epithelial MHCII—and thus loss of direct presentation of injected *Cr* antigens—influenced late CD4 T cell developmental trajectories revealed pathways through which alternate programming of CD4 memory T cells might be mediated (**Fig. 6**). Consistent with our previous findings^1^, deficiency of antigen presentation by colonocytes altered the number of effector CD4 T cells during active infection but had no significant impact on the relative composition of effector subsets (**Fig. 6a,b; Supplementary Table 2,** and **data not shown**). Thus, while there were modest shifts in the relative numbers of Th1 and Th17 cells, for example, these were minor. Conversely, while only modest changes in total memory cell numbers were found at all time points—suggestive of full occupancy of limiting niche space—we found significant shifts in the composition of memory cell subsets: Epithelial-resident Trm cells, both CD103^−^ and CD103^+^, were more dependent on epithelial cell antigen presentation for their development (**Fig. 6a,c** and **data not shown**), hence *H2-Ab1*^Villin^ mice exhibited a shift in memory T cell composition favoring central memory T cells (Tcm) with corresponding decreases in both tissue-resident (Trm) and effector-memory (Tem) populations (**Fig. 6c**). This was supported by flow cytometric findings, which demonstrated significantly increased CD44^+^CD62L^+^ Tcm cells in the mucosa of *H2-Ab1*^Villin^ mice compared to control mice at day 42 (**Fig. 6d-e**). Conversely, both CD69^+^CD103^−^ and CD69^+^CD103^+^ Trm populations were reduced in *H2-Ab1*^Villin^ mice (**Fig. 6f-g**). Thus, the ratio of Tcm to Trm cells was significantly skewed in favor Tcm cells in *H2-Ab1*^Villin^ mice (**Fig. 6h**). In view of our results establishing differential programming of Trm and Tcm cells contingent on injected versus non-injected Cr antigens (**Fig. 4**), these findings indicated that T cell recognition of antigen expressed by DCCs was required for optimal development of Trm cells whereas absence of non-professional antigen presentation favored default to a Tcm fate—and disposition to GALT rather than intraepithelial or lamina propria compartments^60^.

**Fig. 6:**
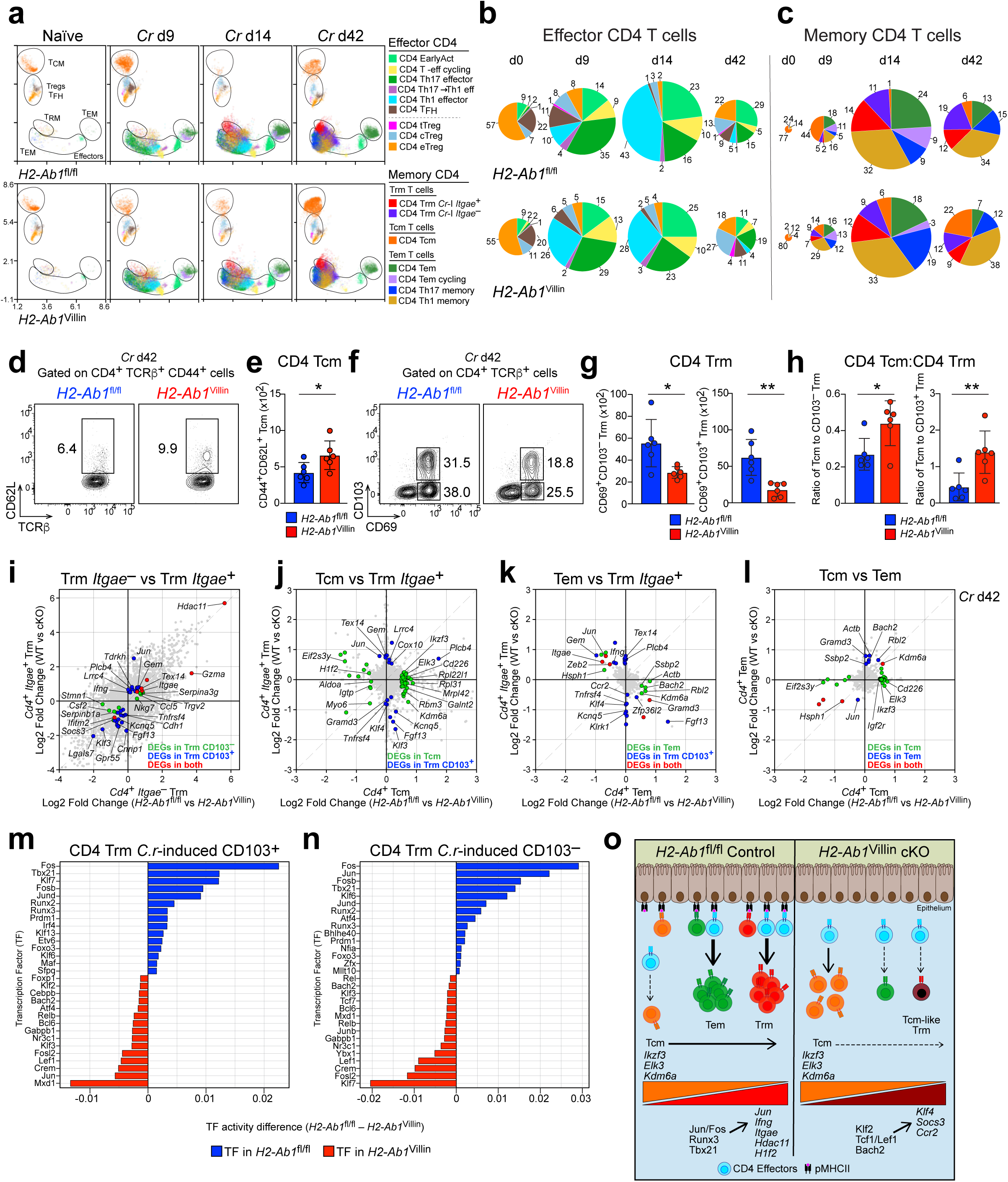
Epithelial cell antigen presentation directs Trm cell transcriptional programming. **a**, UMAP analysis of CD4 effector and memory populations from inidcated *H2-Ab1*^fl/fl^ and *H2-Ab1*^Villin^ mice. **b,c**, Pie charts of the percentages of CD4 effector (**b**) and memory (**c**) clusters from naïve and days 9, 14, and 42-infected *H2-Ab1*^fl/fl^ and *H2-Ab1*^Villin^ mice. **d,e**, Colonic IE cells from day 42-infected *H2-Ab1*^fl/fl^ and *H2-Ab1*^Villin^ mice were stained for CD4, TCRβ, CD44, CD62L, and analyzed by flow cytometry (**d**) to quantify the number of CD44^+^CD62L^+^ Tcm cells (**e**). 3-5 mice per group; *n*=2 independent experiments. Two-tailed unpaired *t*-test; *p≤0.05. Colonic IE cells from day 42-infected *H2-Ab1*^fl/fl^ and *H2-Ab1*^Villin^ mice were stained for CD4, TCRβ, CD69, CD103, and analyzed by flow cytometry (**f**) to quantify the number of CD69^+^CD103^−^ and CD69^+^CD103^+^ Trm cells (**g**). 3-5 mice per group; *n*=2 independent experiments. Two-tailed unpaired *t*-test; *p≤0.05 and **p≤0.01. **h**, Ratio of Tcm cells to CD69^+^CD103^−^ and CD69^+^CD103^+^ Trm cells from the colonic IE compartment in day 42-infected *H2-Ab1*^fl/fl^ and *H2-Ab1*^Villin^ mice. Two-tailed unpaired *t*-test; *p≤0.05 and **p≤0.01. **i**, Two-way scatter plot of DEGs from CD103^−^Trm vs DEGs from CD103^+^ Trm clusters from day 42 scRNA-seq data. Two-way scatter plots of DEGs from Tcm vs DEGs from CD103^+^ Trm clusters (**j**), DEGs from Tem vs DEGs from CD103^+^ Trm clusters (**k**), and DEGs from Tcm vs DEGs from Tem clusters (**l**) from day 42 scRNA-seq data. pyScenic TF regulon analysis performed in CD103^+^ Trm (**m**) and CD103^−^ Trm (**n**) from day 42 scRNA-seq data. **o**, Summary schematic depicting normal effector CD4 T cell development during acute *Cr* infection and subsequent differentiation into memory T cell subsets (Tcm, Tem, and Trm) in *H2-Ab1*^fl/fl^ mice. In *H2-Ab1*^Villin^ mice lacking epithelial MHCII, Tem and Trm populations are reduced with a reciprocal increase in Tcm cells. Additionally, Trm cells that develop in *H2-Ab1*^Villin^ mice display an altered transcriptional program compared to those from *H2-Ab1*^fl/fl^ mice.

Differential gene expression analysis revealed distinct transcriptional programs between memory subsets contingent on epithelial antigen recognition (**Fig. 6i-l**). The greatest dispersion of gene expression alterations was found in Trms (**Fig. 6i-k**). For example, CD103^+^ Trm cells developing in *H2-Ab1*^fl/fl^ mice expressed higher levels of transcripts of tissue residency markers, including *Itgae* (CD103), *Jun*, *Ifng*, *Ccl5*, and *Gzma*. One of the most differentially expressed genes contingent on epithelial MHCII was *Hdac11*, the sole class IV histone deacetylase that has a focused action in repressing effector gene expression in memory T cells^61^, suggesting that this epigenetic modifier may contribute to both CD103^−^ and CD103^+^ Trm cell quiescence in cells developing downstream of IEC presentation of antigen. In agreement with a report that examined small intestinal Trm cells^62^, expression of TF transcripts of *Hic1* and *Ahr* were enriched in both CD4 and CD8 Trms induced by *Cr* infection, but were not contingent on IEC antigen presentation, indicating that other factors control up-regulation of these factors in developing Trm cells. Similarly, transcripts for *Nr4a1* (Nur77), which was particularly enriched in CD103^−^ CD4 Trms, was not differentially expressed between genotypes. Further, in *H2-Ab1*^Villin^ mice, the transcriptional identity of Trm cells that did develop was altered, with increased expression of *Klf3*, *Klf4*, *Socs3*, and *Ccr2*—markers typically associated with central memory or migratory programs^63–65^. These cells retained some tissue residency markers such as *Itgae* and *Cd69* but lost the effector-associated genes characteristic of *H2-Ab1*^fl/fl^ Trm cells. Notably, although the relative cell numbers of Tcm diverged between *H2-Ab1*^fl/fl^ and *H2-Ab1*^Villin^ mice, few genes were differentially expressed, consistent with similar “default” programming in memory cells that develop in absence of antigen recognition on IECs.

Transcription factor regulon analysis of CD4 T cells from *H2-Ab1*^fl/fl^ versus *H2-Ab1*^Villin^ mice further supported these findings, showing that both CD103^+^ and CD103^−^ Trm cells developing in *H2-Ab1*^fl/fl^ mice exhibited higher activity of Fos, Tbx21, Jun, Runx2, Runx3 and Prdm1—factors associated with more terminal development, tissue residency and sustained effector competency^66–68^ (**Fig. 6m-n**). In contrast, Trm cells that developed in *H2-Ab1*^Villin^ mice showed increased activity of Mxd1, Tcf7, Bcl6 and Bach2—factors that typically promote more plastic, reversible transcriptional programming and suppression of tissue residency that are features of central memory cell fate. Klf2, Tcf7/Lef1, and Bach2 have been shown to specifically regulate the expression of *Klf4*, *Socs3*, and *Ccr2*, differentially expressed gene signatures noted above. Interestingly, the Klf7 regulon was a prominent feature of both CD103^+^ Trm cells that developed in *H2-Ab1*^fl/fl^ mice and CD103^−^ Trm cells from *H2-Ab1*^Villin^ mice, indicating that its role in promoting adaptation for memory cell survival and prevention of exhaustion was uncoupled from EC antigen presentation. Together, these data are consistent with the less restricted transcriptional identity of Tcm cells compared to the more committed phenotype of Trm cells and reveal that MHCII-dependent antigen presentation by DCCs provides critical signals that shape the transcriptional landscape of developing memory T cells—actively promoting tissue-resident memory cell programming to avoid default central memory programming.

Collectively, these results support a model whereby translocation of bacterial antigens into intestinal epithelial cells during *Cr* infection enables critical programming of antigen-specific CD4 T cells through epithelial MHCII presentation (**Fig. 6o**). Epithelial presentation of injected antigens not only amplifies the magnitude of the T cell response, resulting in many more effectors in the involved mucosa during active bacterial infection, but also directly influences memory T cell fate decisions, promoting the development and maintenance of tissue-resident memory populations that provide frontline defense at the intestinal barrier. The persistence of epithelial MHCII expression and continued epithelial-T cell interactions after pathogen clearance suggests that epithelial cells serve as both initiators and long-term maintainers of barrier immunity.

## Discussion

In this study, we have discovered that all bacterial antigens are not immunologically equal. At least for bacteria that use a T3SS to colonize host barrier epithelia, there is a hierarchy in the induction and mucosal recruitment of CD4 T cells contingent on the distribution of bacterial antigens, strongly favoring those delivered to the cytosol of the host epithelium—independent of epitope binding affinity for MHCII or heterogeneity of TCR avidity for MHC-peptide complexes generated. Indeed, antigen localization eclipsed antigen load as the principal determinant of the magnitude and developmental trajectory of the T cell response, and injected bacterial effectors were the dominant class of antigen that programmed development of Trm cells, ensuring that Trm responses were focused on epithelium-associated antigens. Coupled with our recent report describing a role for epithelial cell antigen presentation to efficiently recruit CD4 T cell help for barrier defense during active infection^1^, the current findings consolidate the importance of these non-professional APCs in guiding the adaptive immune response during and following a pathogenic challenge.

The route by which antigens enter the host has long been appreciated as an important determinant of immunogenicity. This concerns both the pathways through which antigens traverse barrier epithelia and by which they are routed into, and within, antigen-presenting cells. Antigen form is similarly important. Early studies highlighted differences in immunogenicity of particulate versus soluble antigens^69,70^, with particulate antigens particularly suited to uptake by phagocytic APCs (i.e., DCs and activated macrophages), which deploy an array of scavenger receptors that direct extracellular material to endolysosomal compartments for processing and loading onto MHCII (MIICs)^71^. Soluble antigens are poorly taken up by phagocytic APCs but can be efficiently delivered to MIICs of B cells if harboring repeated epitopes that are bound by surface immunoglobulin^72^. In contrast to these professional APCs, epithelial cells lack efficient mechanisms for uptake of pathogen-associated antigens and therefore rely on the bacterium itself to deliver antigens for presentation. Unlike invasive bacteria such as pathogenic *Salmonella enterica* serovars or *Listeria monocytogenes*, which activate mechanisms for direct bacterial entry into intestinal epithelial cells, A/E pathogens such as *Cr* lack such mechanisms and only bacterial antigens delivered by the T3SS enter host IECs. In this sense, bacterial “invasion” is a relative term: Although *Cr* does not penetrate cECs as an intact cell, its injection of effector molecules into these cells represents a barrier breach that is clearly detected by T cells primed against these antigens.

As demonstrated herein, the quantity and quality of the clonal CD4 T cell response were highly affected by the disposition of bacterial antigens: bacterium- vs epithelium-associated, and, within the epithelial cell, membrane- vs cytosol-associated. The superior immunogenicity of injected cytosolic *Cr* antigens indicated that diverse other routes of antigen uptake in the intestines—capture of lumenal Ags by transepithelial dendrites (TEDs) projected by CX3CR1 macrophages^73^, bulk and receptor mediated transport across M cells^74^, uptake via Goblet cell-associated passages (GAPs)^75^ or antibody-mediated transport via the neonatal Fc receptor (FcRN)^76^, among others—were less contributory. The relatively weak response to bacterially-associated antigen was unexpected, as antigens of this class, such as intimin, are targets of an IgG antibody response that is important in thwarting *Cr* pathogenicity and clearing infection^33^. While for intimin-gp66 this might be explained by its relatively low antigen dose, the dose of naturally expressed espZ-gp66 was similar—despite inducing a much more potent response—and dose alone could not explain the similarly muted response to espZ-Δ1-21, which was expressed at comparable levels to the injected full-length variant yet induced greater than 100-fold fewer SMARTA T cells. This suggests that for A/E pathogens at least, dendritic cells are more efficient at taking up epithelial cell-associated antigen, whether captured through direct uptake of debris from damaged or dying IECs or via transfer through an intermediate, such as CX3CR1^+^ mononuclear phagocytes as occurs for GAP-mediated uptake^77^. Why this might be so—and why the cytosolic antigens espZ and nleA would be preferentially taken up over tir—is unclear but may reflect differential display of scavenger receptor targets by the damaged basolateral versus apical membrane domains of IECs, or simply a propensity of the apical membrane to be extruded into the intestinal lumen upon injury of infected IECs. This will require further study.

Because injected *Cr* antigens bypass the classical pathway of MHCII loading, our results favor a non-classical pathway involving one of the autophagy-mediated mechanisms (macroautophagy, microautophagy or chaperone-mediated autophagy) for loading injected cytosolic antigens into compartments for processing and loading onto MHCII proteins (MIICs)^78–80^. And since espZ and nleA lack KFERQ-like motifs it is unlikely they are processed by the chaperone-mediated autophagy (CMA) pathway. Moreover, there was clear stratification of the clonal response to the three injected *Cr* proteins, with espZ-gp66 and nleA-gp66 demonstrating more potent induction of the SMARTA response than tir-gp66, despite the latter showing the greatest density of IEC-associated antigen. This is consistent with greater entry of cytosolic rather than integral membrane proteins into the autophagy pathway, and, coupled with the absence of a targeting sequence, is most characteristic of macroautophagy, although this, too, will require further study.

The mechanisms controlling the development of different types of T cell memory are an area of active investigation^81,82^. Our findings highlight an important role for CD4 T cell recognition of MHCII-peptide complexes displayed on infected epithelial cells in programming tissue-residency. In view of the strong bias towards cytosolic antigens for IEC presentation, this implies a central role for CD4 Trm cell programming by injected bacterial proteins—as demonstrated by the striking bifurcation in specification of Trm versus Tcm cells driven by injected cytosolic or bacterially associated proteins, respectively. This makes biological sense, as adaptive immunity has evolved to focus immune resources where needed; linking antigen recognition on IECs to induction of local tissue residency ensures deployment of Trm cells to the site they will encounter antigen anew upon reinfection. Notably, while it has been suggested that the intestinal Trm pool spans both intraepithelial and lamina propria (LP) mucosal compartments, our findings indicate that relatively few memory T cells populate LP, indicating instead that the major sites of memory cell accumulation are within the epithelium and GALT. The preferential positioning of Trm cells to the epithelium also makes sense, as our scRNA-seq data establish that the T cell-sustaining cytokines, IL-7 and IL-15, are differentially produced by different populations of colonocytes, with IL-15 produced by mature IECs near the luminal surface and IL-7 by immature cIECs located deeper in crypts. This aligns well with recent spatial transcriptomics data describing two subsets of CD8 Trm cells arrayed along the crypt-villus axis of small intestine in proximity to spatially segregated epithelial IL-7 or IL-15 producers^83^. Although new spatial transcriptomics studies will be needed to resolve the transcriptional signature of the few T cells found in LP in the memory phase, we suspect they are effector-memory (Tem) cells that are transiting the mucosa in their migration between blood, tissue and lymphatics^84,85^.

Our scRNA-seq data provide a wealth of data on the bidirectional information exchange that occurs between cECs and CD4 T cells, exposing transcriptional networks underpinning altered cEC programming downstream of T cell help and reciprocal programming of T cells that engage antigens presented by cECs. Dominant signaling into cECs was downstream of IFNγ, reflecting the greater number of Th1 cells present at the peak of effector T cell response. IFNγ is not only critical for programming important barrier defense functions in the epithelium^5^, it is also critical to activate the antigen processing and presentation machinery required to recruit Ag-dependent help for the epithelium, whether downstream of IFNγ itself—or IL-22, the other major epithelium-protective cytokine that relies on EC antigen presentation to prevent lethality by *Cr*^1,86–88^. The findings herein reveal heretofore unknown actions of IFNγ signaling in IECs that will be a basis of future studies. It should be noted that the progressive shift in the Th17/Th22–Th1 cell balance resulting from transdifferentiation of Th17 cell precursors may be linked to Trm specification, since the transcriptional programming of Trm cells is more aligned with the Th1 program, many features of which are shared with CD8 Trm cells. Thus, primed T cells that enter the infected mucosa as members of the more plastic Th17 program would appear to be progressively constrained as they transition to Th1 and Trm cells, which lose stem-like characteristics as they develop^89,90^.

Integration of the scRNA-seq data analysis with other findings in our study further elucidate features of the T cell response to antigens displayed on cECs. Although there were substantial shifts in relative numbers of Trm and non-Trm memory cells contingent on epithelial antigen presentation, a requirement for Trm specification by EC-presented antigen was not absolute. However, Trm cells that developed in MHCII-deficient mice did retain some transcriptional signatures of Tcm cells, suggesting that signals promoting full Trm cell commitment (e.g., TGFβ^59,91^) may be deficient without sustained engagement on antigen-bearing ECs. Thus, our findings suggest that T cell recognition of EC-presented antigen may not deliver a unique set of signals without which Trm development cannot occur, rather, sustained engagement of T cells in the epithelial microenvironment drives to completion antigen-dependent programming associated with extinction of a more plastic, migratory memory lifestyle, and commitment to survival maintained by local epithelial factors, such as IL-15 and IL-7. Also, the balance of CD103^−^ and CD103^+^ Trm cell numbers and their transcriptional signatures were not appreciably affected by loss of EC antigen presentation, and did not appear to reflect absence of close interactions with epithelial APCs, as labeling with uLIPSTIC system established that most Trm cells of both subsets actively engaged ECs during the memory phase, albeit more limited for the CD103^−^ subset. Whether this reflects dwell time differences, positioning differences along the crypt-surface axis within the epithelium, or other factors, warrants further study. Finally, findings herein extend characterization of transcriptional networks activated to program tissue residency in CD4 T cells, identifying remarkable similarity to those of better-characterized CD8 Trm cells, but also exposing new players, detailed functional contributions of which await future studies.

In summary, our findings indicate that MHCII-dependent interactions between cECs and CD4 T cells are bidirectional and preferentially mediated by antigenic peptides derived from bacterial proteins injected into the cytosol of host cECs. This supports a model wherein the processing and presentation of these antigens by cECs both elicit T cell help crucial for acute barrier protection and promote development of epithelium-resident memory T cells that mediate an accelerated protective response upon repeat pathogen challenge. Further, these findings support a model of T cell programming wherein the antigen-presenting EC does not provide indispensable signals for memory cell subset development, rather the recruitment of CD4 T cells to sustained interactions with the epithelium afforded by EC antigen presentation increases exposure of the T cells to a microenvironment enriched for Trm-promoting signals, such as TGFβ. Absent these interactions, which are predicated on recognition of injected bacterial antigens that load the epithelial APP pathway, the T cell fails to progress in memory programming and exits the infected mucosa to reenter the circulating memory pool. These findings have important implications for mucosal vaccine design, suggesting that targeting antigens to the cytosol of epithelial cells may enhance protective Trm responses at barrier tissues such as the intestines. They may also give rise to new therapeutic strategies that target pathological immune memory responses at barrier tissue sites.

## Methods

Detailed methods are included in the Supplementary Information Section.

## Acknowledgements

The authors thank B. A. Vallance, W. Deng, J. Moon, C. L. Weaver and members of the Weaver laboratory for helpful discussions; H. Turner, B. Dale, K. Cantrell, C. Shah, and S. Shah for assistance with animal husbandry; V. S. Hanumanthu, H.C. Pal, C.-W. Sun, S. Liu, M. Basu, C. Iradukunda, K. Stortz, T. Houser, and H. Johnson at the UAB Flow Cytometry and Single Cell Core Facility; S. Williams and R. Grabski at the UAB High Resolution Imaging Facility (HRIF); and D. Mohr at the Genetic Resources Core Facility (GRCF) at Johns Hopkins University. This work was supported by R01 grant funds from NIH/NIAID (C.T.W.) and NIH T32 and F30 support (C.G.W.).

## Author Contributions

C.G.W., C.L.Z., and C.T.W. conceived the project and wrote the manuscript. C.G.W., C.L.Z., M.P.A. and C.T.W. performed the experiments and/or interpreted the results. L.W.D. and J.R.S. assisted with development of epitope-tagged *Cr* strains. L.K. and M.G. performed the LSFM and analysis. N.T.-A., Y.W., and J.M. performed the smFISH and analysis. H.S. and C.X. generated Western blot data. S.N.H. and Y.N.-K. conducted *in vitro* fertilization procedures to maintain all mouse lines with the same C57BL/6 microbiota. R.D.H. assisted with scRNA-seq experiments. M.P.A. and V.O. performed bioinformatics analyses of scRNA-seq data. B.F.F. provided technical expertise and contributed to data interpretation.

## Supplementary Information

## Methods

### Mice

C57BL/6 (WT; stock #000664), C3H/HeJ (stock #000659), *H2-Ab1* floxed (stock #013181)^1^, SMARTA-1 CD45.1 (stock #030450)^2^, *Villin*-cre (stock #021504)^3^, and *Villin*-cre/ERT2 (stock #020282)^4^ mice were purchased from Jackson Laboratory. Rosa26^uLIPSTIC5^ mice were a gift from Gabriel D. Victora. MHCII-EGFP^6^ mice were a gift from Hidde L. Ploegh. *H2-Ab1*^fl/fl^ and *H2-Ab1*^Villin^ mice were screened by PCR and flow cytometry to exclude mice with spontaneous, Cre-independent germline deletion of MHCII, as previously reported^7^. All mice undergo rigorous screening to exclude SFB, *Helicobacter* spp., and other potential pathobionts. All mouse strains were bred and maintained at UAB in accordance with IACUC guidelines.

### Bacterial strains and infections

*Citrobacter rodentium (Cr)* strain, DBS100 (ATCC) was used for scRNA-seq experiments. For flow cytometry analysis and cytospin, we used a strain of *Cr* expressing GFP^8^ (derived from DBS100; kindly provided by Bruce A. Vallance). A fresh, single colony was grown in 10 mL LB at 37°C with rotation at 225 rpm for 12-14 hrs. Next day, 1 mL of overnight culture was added to 250 mL LB broth (Fisher), incubated at 37°C with rotation (225 rpm) for 3-3.5 hrs until OD_600_ reached 1.0 on spectrophotometer (ThermoFisher Spectronic 200). Bacteria were pelleted at 25°C, 3000 rpm for 15 minutes and then resuspended in 5 mL sterile 1× PBS. Mice were inoculated with 2 × 10^9^ CFU in a total volume of 100 µl of PBS by gastric gavage.

### Generation of epitope-tagged *C. rodentium* strains

We previously reported the detailed methods for this process^7^. Briefly, DNA constructs were synthesized by GeneArt (ThermoFisher) and then cloned into the pCAL52^9^ shuttle vector (kindly provided by Sebastian E. Winter). Transformed EcNR1Δasd *E. coli*^10^ (kindly provided by Michael J. Gray) containing the cloned pCAL52 plasmid was grown O/N in LB broth with 25 mg/mL chloramphenicol and 1 mg/mL 2,6-diaminopimelic acid (DAP, Sigma) and *Cr* (DBS100 or *Cr*-GFP) was grown O/N in LB broth. Puddle mating was performed on an LB agar plate containing 1 mg/mL DAP and incubated at 30°C O/N to promote conjugation. The puddle mating was then plated on LB agar plates containing 25 mg/mL chloramphenicol (without DAP) and incubated at 37°C O/N. Single-crossover merodiploid clones were confirmed by PCR and then selected in 25% sucrose LB broth for 3-4 days with daily plating on LB agar containing 40 mg/mL 5-Bromo-4-chloro-3-indolyl phosphate p-toluidine salt (BCIP, Sigma) for blue-white detection of clones that had successfully undergone a double-crossover event to remove the *phoN*-containing pCAL52 backbone. White colonies were confirmed as either WT revertants or homozygous insertion mutants by PCR using the following forward and reverse primers:

eae_F 5’-GATGGCGTATTCTGAGGCAG-3’

eae_R 5’-CGGGTGCCGGGTTTATATGAT-3’

tir_F 5’-GAAGAGCCTATTTATGATGAAGTCGC-3’

tir_R 5’-GTTCTCTGAAACATTGACATACTCC-3’

nleA_F 5’-AGTGTGTCGTGGTCTTGCAA-3’

nleA_R 5’-TATGGTGGGCAGGCGCGACA-3’

espZ_F 5’-CGGGAATTGCAGCAATGTGT-3’

espZ_R 5’-GGTTGGGGCTAACGGAGTAT-3’

xylE_F 5’-AGGGTTCACCAGCATCAACC-3’

xylE_R 5’-TCTTACGTGCCGATCAACGT-3’

### Western blot detection of epitope-tagged bacterial proteins

Sorted *Cr*-GFP-bound EpCAM^+^ IECs, *Cr* pellets from culture in DMEM, or supernatant proteins from *Cr* culture in DMEM concentrated with trichloroacetic acid (TCA) precipitation (O/N at 4°C) were resuspended in 2× Laemmli buffer (4% SDS, 20% glycerol, 0.005% bromophenol blue, 0.125 M Tris-HCl (pH 6.8), and 10% b-mercaptoethanol). Lysates were transferred to 1.5 ml Eppendorf tubes and briefly sonicated. Total proteins were loaded onto 8-15% SDS-PAGE gel for electrophoresis separation. After electrophoresis, proteins on the gel were transferred onto 0.45 µm PVDF membrane (Millipore, #IPVH00005) in a sponge sandwich. Membranes were blocked with 5% non-fat milk (Bio-Rad, #170-6404) and probed with primary antibodies overnight on a shaker at 4°C. Membranes were then washed and incubated with HRP-conjugated secondary antibodies for 1 hour at room temperature. The membranes were then incubated with Immobilon Crescendo Western HRP substrate (Millipore, WBLUR0500) for 2-5 min before imaging with CL-X Posure X-Ray film (ThermoFisher). β-actin was used as a loading control for IECs and GFP or DNAK were used as a loading controls for *Cr*. The following antibodies were used: mouse anti-HA (6E2, Cell Signaling Technology), rat anti-HA (3F10, Sigma), mouse anti-FLAG (M2, Sigma), mouse anti-b-actin HRP (sc-47778, Santa Cruz Biotechnology), anti-GFP (rabbit polyclonal, ThermoFisher), mouse anti-DNAK (8E2/2, Enzo), anti-rabbit HRP (goat polyclonal, ThermoFisher), anti-rat HRP (goat polyclonal, ThermoFisher), and anti-mouse HRP (goat polyclonal, ThermoFisher).

### IEC and intraepithelial lymphocyte (IEL) isolation

Intestinal tissue was flushed with 1× PBS, cut into 4 cm mid-distal colon, opened longitudinally and then cut into strips of 1 cm length. Tissue pieces were incubated for 20 min at 37°C with 1 mM DTT (Sigma), followed by 2 mM EDTA (Invitrogen) in H5H media (1× HBSS (Fisher), 5% FBS, 20 mM HEPES (Fisher), and 2.5 mM 2-β-ME (Gibco)). Tissue pieces were vortexed briefly after each 20 min incubation, followed by washing with H5H prior to centrifugation at 1750 rpm for 10 min at 4°C. IECs were then purified on a 40%/75% Percoll (Cytiva) gradient by centrifugation for 20 min at 25°C and 2000 rpm with no brake. For analysis of *Cr*-GFP attached to IECs, tissue pieces from 4 cm of mid-distal colon were incubated for 20 min at 37°C with 1 mM EDTA in H5H media, followed by gentle mixing and washing with H5H.

### Lamina propria lymphocyte (LPL) isolation

Colons were flushed with 1× PBS, opened longitudinally then cut into small pieces and placed in H5H media. Tissue was then minced in 1.5 mL microcentrifuge tubes for 2-3 min before being transferred into scintillation vials with 10 mL of complete R10 media (1× RPMI 1640 (Corning), 10% FBS, 1× Pen/Strep (Corning), 1× NEAA (Corning), 1mM Sodium pyruvate (Corning), 2 mM L-glutamine (Corning), and 2.5 mM 2-β-ME (Gibco)) with collagenase IV (Sigma, 100 U/mL) and DNase I (Sigma, 20 mg/mL). Tissue was digested at 37°C for 40 min with stirring followed by filtering over a 70 mm filter and washing with complete R10 media. Cells were then centrifuged at 1750 rpm at 4°C for 10 min and then purified on a 40%/75% Percoll as described above. Where indicated, cells were stimulated with phorbol-12-myristate-13-acetate (PMA, Sigma, 50 ng/mL) and ionomycin (Sigma, 750 ng/mL) at 37°C for 3-4 hrs in the presence of GolgiPlug (BD Biosciences).

### Cytospin and immunofluorescence staining of bacteria-bound IECs

Sorted epithelial cells (2–5×10^5^ cells/mL) were cytospun onto glass slides at 1,500 rpm for 2 min at room temperature. Cells were then dried for 10 min, fixed with 4% RT paraformaldehyde (Electron Microscopy Sciences) for 10 min and then permeabilized with 0.1% Triton X-100 (Sigma) for 10 minutes prior to staining. The following antibodies and reagents were used: anti-GFP (A-11122; ThermoFisher), anti-goat IgG (ThermoFisher), anti-HA tag (polyclonal; Novus Biologicals), anti-rabbit IgG (ThermoFisher), and ProLong Diamond Antifade Mountant with DAPI (ThermoFisher).

### Flow cytometry and cell sorting

Colon cells were stained with Fc Block (anti-CD16/32, clone 2.4G2) followed by staining with fluorescent-labeled antibodies in IEC buffer (1× PBS with 5% FBS and 2mM EDTA to reduce cell clumping) for IECs and IELs or 2% FBS in 1× PBS for LPL cells on ice in 1.5 mL microcentrifuge tubes or in a 96-well plate. For intracellular staining, cells were fixed and permeabilized using BD Cytofix/Cytoperm kit (BD Biosciences). Samples were acquired on an Attune NxT flow cytometer (Life Technologies) or FACSymphony A3 (BD Biosciences) and analyzed with FlowJo software. Cells were sorted on a FACSymphony S6 (BD Biosciences). The following antibodies/reagents were used: anti-CD45 (30-F11; Biolegend), anti-EpCAM (G8.8; ThermoFisher), anti-MHCII (M5/114.15.2; Fisher), Live/Dead Fixable Near-IR dead cell dye (ThermoFisher), anti-CD4 (RM4-5; BioLegend), anti-TCRb (H57-597; ThermoFisher), anti-CD44 (IM7; BioLegend), anti-CD45.1 (A20; ThermoFisher), anti-CD45.2 (104; BD Biosciences), anti-CD69 (H1.2F3; ThermoFisher), anti-CD103 (2E7; ThermoFisher), anti-CD19 (1D3/CD19; Biolegend), anti-CD11b (M1/70; Biolegend), anti-CD11c (N418; Biolegend), anti-CD62L (MEL-14; Biolegend), anti-PD-1 (29F.1A12; Biolegend), anti-CXCR5 (L138D7; Biolegend), anti-IL-17A (TC11-18H10; BD Biosciences), anti-IFNg (XMG1.2; ThermoFisher), and anti-IL-22 (1H8PWSR; ThermoFisher).

### Quantitative RT-PCR (RT-qPCR)

cDNA synthesis was performed with iScript Reverse Transcription (RT) Supermix (Bio-Rad) according to manufacturer’s instructions. cDNA amplification was performed with SsoAdvanced Universal SYBR Green Supermix (Bio-Rad) in a Biorad CFX qPCR instrument. The following primer sequences were used:

Gapdh_F 5’-TCCATGACAACTTTGGCATTG-3’

Gapdh_R 5’-CAGTCTGGGTGGCAGTGA-3’

Il7_F 5’-CAGTATCACAAGGCACACAAAC-3’

Il7_R 5’-CTCTCAGTAGTCTCTTTAGGAAACAT-3’

Il15_F 5’-ATCCATCTCGTGCTACTTGTG-3’

Il15_R 5’-CTATCCAGTTGGCCTCTGTTT-3’

### Adoptive T cell transfer

Approximately 20-24 hours prior to adoptive transfer of CD45.1^+^ SMARTA cells mice were administered 0.2 mg/kg anti-CD4 (BioXcell, GK1.5) in 1× PBS i.p. Spleens and lymph nodes from CD45.1^+^ SMARTA donors were collected and dissociated in complete R10 media prior to centrifugation at 1500 rpm for 5 min at 4°C. RBCs were removed from the cell suspension by ammonium-chloride-potassium (ACK, Quality Biological Inc.) lysis for 2 mins prior to quenching with complete R10 media. Naïve CD4^+^ T cells from pooled spleens and lymph nodes were purified using the naïve CD4^+^ mouse T cell Isolation kit per manufacturer’s instructions (Miltenyi Biotec). Purified T cells were transferred by i.v. retro-orbital injection (1 × 10^5^ total cells) into recipients, followed by *Cr* infection on the same day.

### Light sheet microscopy and 3D image analysis

Mice received 20 mg anti-EpCAM-AF647 (G8.8, Biolegend) 24 hours and 15 mg anti-CD45.1-AF594 (A20, Biolegend) 3 hours prior to sacrifice and tissue collection. Mid-distal colons were collected into ice-cold 4% PFA (Electron Microscopy Sciences) and stored at 4°C O/N before subsequent washes in ice-cold 1× PBS. Tissue was then dehydrated in increasing concentrations of ethanol before clearing in 100% ethyl cinnamate (ECi, Sigma). Tissue was stored in 100% ethyl cinnamate at RT until imaging. Whole-organ LSFM of optically cleared middle-distal and distal colon samples was performed on an UltraMicroscope Blaze (Miltenyi Biotec, Germany) in multicolor acquisition mode as previously shown^11^. The microscope system employs the ImSpector software (LaVision Biotec, Germany). Samples were positioned on a steel holder within the imaging chamber, which is filled with 100% ethyl cinnamate. Tissue orientation was stabilized using UV-curable optical adhesive (Norland Products). The imaging was performed with a total magnification of 4× (4× detection objective, 0.35 detection NA; 1.0 zoom), bi-directional light-sheet illumination, and an exposure time of 100 ms. The light-sheet thickness was set to 3.9 µm, and the sheet width was adjusted to 60-70%, corresponding to ∼12-14 mm effective illumination field width. The entire intestinal tube – covering a total z-range of approximately 1700-2600 µm per sample – was imaged with a z-step size of 2 µm between images. CD45.1^+^ SMARTA cells, labeled with anti-CD45.1-AF594, were excited using a 595/20 nm OPSL (50 mW) and detected with a 630/30 nm bandpass filter. Epithelium, stained with anti-EpCAM-AF 647, was excited with a 630/30 nm OPSL and detected by a 680/30 nm bandpass filter. 3D rendering and quantitative image analysis were performed in IMARIS 10.1.0 (Oxford Instruments, UK). Raw LSFM datasets were converted into IMARIS format using the IMARIS File Converter. Quantification of total CD45.1^+^ SMARTA cell counts and subsequent classification into epithelial, lamina propria and gut-associated lymphoid tissue (GALT) compartments was conducted using the IMARIS Surface module. For both imaged fluorescence channels, iterative rounds of machine-learning-assisted threshold refinement (IMARIS machine-learning classifier) were performed to ensure high-accuracy surface segmentation while excluding non-specific autofluorescent structures. First, an epithelial surface was generated based on the fluorescent intensity of the EpCAM channel to define anatomical boundaries for the relative distribution of CD45.1^+^ SMARTA cells in the colon. Epithelial CD45.1^+^ cells were defined as surfaces with a distance to the EpCAM surface <0 µm, whereas lamina propria CD45.1^+^ cells were defined by a distance >0 µm. The third category, GALT-associated CD45.1^+^ cells, was manually assigned based on the distinctive morphological appearance of colonic patches and isolated lymphoid follicles (ILFs). Throughout the entire rendering workflow, potential non-specific or autofluorescent structures were carefully inspected in all anatomical planes and removed manually when necessary.

### smFISH staining of infected and uninfected cecal sections

Samples were prepared and imaged using published smFISH protocols^12^, with minor modifications. Briefly, cecal tissues were fixed in methacarn solution (60% methanol, 30% chloroform, and 10% glacial acetic acid; v/v/v) at 4°C for 48 hours to preserve both tissue morphology and RNA integrity. Fixed specimens were subsequently washed with 100% methanol (Sigma, MX0480) for 35 minutes at 4°C for a total of two times. Next, the tissue was washed in 100% ethanol (ThermoFisher, 04-355-223) for 30 minutes at 4°C for a total of two times to eliminate residual fixative. The tissues were then stored in fresh 100% ethanol at 4°C for 48-96 hours prior to paraffin embedding, which was performed by the Rodent Histopathology Core at the Dana-Farber/Harvard Cancer Center. During the embedding process, tissues were incubated in xylene (VWR, 89370-088) for 1.5 hours at room temperature, repeated twice, followed by infiltration with molten paraffin wax (Leica, 3801340) for 4-5 hours at 60 °C. Samples were subsequently embedded in paraffin blocks (Leica, 3801320). Embedded tissues were sectioned at a thickness of 4 µm and mounted on poly-L-lysine-treated, silanized coverslips to ensure flat adhesion during downstream staining procedures. Sections were baked at 60°C for 20 minutes, then washed four times with xylene (Sigma, 534056) for 2.5 minutes each to remove paraffin. Residual xylene was cleared by two washes in 100% ethanol for 3 minutes, followed by rehydration through sequential 95% and 70% ethanol washes for 1 minute each. Rehydrated sections were post-fixed in 4% (v/v) paraformaldehyde (Electron Microscopy Sciences, 15714) in 1× PBS for 10 minutes to crosslink RNA, then washed three times in 1× PBS to remove the PFA. Samples were equilibrated in a 30% formamide solution (deionized formamide, Thermo, AM9344, in 2× SSC) before staining. The probe sequences used in smFISH experiments were designed to hybridize to 16S rRNAs in bacteria and included a 20-nt readout sequence (underlined) appended to each target sequence to allow specific binding of fluorescently labeled oligonucleotides for detection during fluorescence microscopy imaging. The applied *Citrobacter rodentium*-specific probe (5′-CCAGATGGGATTAGCTTGTTGGTGAGGTAA A <u>ACTCCACTACTACTCACTCT</u>-3′), eubacterial probe (5′-<u>AATCTCCTCCAACACTTCTA</u> A CCAGACTCCTACGGGAGGCAGCAGTAGGGA-3′), and corresponding fluorescently-labeled probes (*C. rodentium*: 5’-Alexa750-S-S-AGAGTGAGTAGTAGTGGAGT-3’; Eubacteria: 5’-Cy5-S-S-TAGAAGTGTTGGAGGAGATT-3’) were synthesized and experimentally validated to ensure binding and specificity as previously described^13,14^. Following staining, sections were embedded in a 4% (v/v) polyacrylamide gel prepared from a 19:1 bis-acrylamide mixture (Bio-Rad, 1610144). The gel solution contained 0.05 M Tris-HCl, pH 7.4 (ThermoFisher, 15568-025), 0.3 M NaCl (ThermoFisher, AM9759), 0.03% (w/v) ammonium persulfate (Sigma, 215589), and 0.15% (v/v) N′-tetramethylethylenediamine (TEMED; Sigma, T7024). Embedded sections were cleared overnight at 37 °C using a digestion buffer composed of 2% (v/v) SDS (ThermoFisher, AM9822), 8 U/mL proteinase K (New England Biolabs, P8107S), and 0.25% (v/v) Triton X-100 (Sigma, T8787) in 2× SSC. Cleared samples were washed with 2× SSC for 30 minutes at room temperature a total of five times and stored at 4°C in 2× SSC supplemented with murine RNase inhibitor (1:400 dilution; New England Biolabs, M0314L) until imaging.

### Microscopy imaging and data analysis

Stained samples were imaged on a custom-built Nikon Ti-2 epifluorescence microscope equipped with a Celesta laser light engine (Lumencor, 90-10521), a 60× CFI PlanApo oil objective, a 10× CFI PlanApo air objective, and two Hamamatsu ORCA Flash 4.0 CMOS cameras connected via a TwinCam color splitter (Cairn), as previously described^12^. Samples were mounted in a Bioptechs FCS2 closed-flow chamber (1 mm gasket, Bioptechs 1907-1422-1000) connected to a custom automated fluidics system composed of a peristaltic pump and Hamilton MVP valves. The flow system was programmatically controlled through an established computational pipeline (https://github.com/ZhuangLab/storm-control), allowing automated buffer exchanges and multiple smFISH imaging rounds to be performed sequentially. During imaging, mounted samples were stained for at least 15 minutes with fluorescently labeled oligonucleotide sequences that hybridized to the 20-nt readout regions appended to each rRNA probe. Staining was performed either at the bench or directly on the microscope by configuring the fluidics system to slowly flow a hybridization solution containing the fluorescently labeled oligonucleotides over the sample, ensuring stable probe-target binding. To minimize photobleaching and enhance fluorescence signal intensity, samples were suspended in an oxygen-scavenging imaging buffer containing 4 μM Trolox-quinone, 0.5 mg/mL Trolox (Abcam, AB120747), 1:500 recombinant protocatechuate 3,4-dioxygenase (rPCO; OYC Americas, 46852004), and 5 mM protocatechuic acid (Sigma, 37580) in 2× SSC. During the first round of imaging, samples were bench-top stained with fluorescent probes targeting *C. rodentium* rRNAs and DAPI for 15-20 minutes. The samples were rinsed briefly with 2× SSC to remove unbound dyes, mounted on the microscope, and suspended in imaging buffer prior to being illuminated at 750 nm, 635 nm, and 405 nm to visualize *C. rodentium* rRNA signals, background autofluorescence, and DAPI-stained host structures, respectively. To obtain a tissue-wide image of bacteria localization within host boundaries, multiple fields-of-view are photographed from a given sample using automated stage scanning. After the first imaging round, fluorescent signals were removed by incubating the sample in a cleavage buffer containing 50 mM Tris(2-carboxyethyl)phosphine (TCEP; GoldBio, TCEP25) in 2× SSC. Samples were then washed with fresh 2× SSC to eliminate residual cleavage buffer and cleaved fluorescent molecules. Subsequently, a new set of fluorescently labeled probes was introduced, allowing binding of fluorescently labeled probes to their intended targets, and visualization of eubacterial signals marking the distribution of bacterial populations within the tissue. The produced collection of images from both rounds of microscopy imaging were subsequently stitched together to form a composite mosaic image and displayed using FIJI and Python scripts.

### Single-cell isolation and library preparation

IECs, IELs and LPLs were isolated from the mid–distal colon of *H2-Ab1*^fl/fl^ and *H2-Ab1*^Villin^ mice at four timepoints: naïve (day 0), day 9, day 14, and day 42 post *C.r* infection. For each condition, cells were pooled from 3-4 mice. Following cell isolations as described above, single-cell suspensions were stained and sorted directly into 5 μL PBS supplemented with 5% FBS and 0.1 mM EDTA to obtain live EpCAM⁺ IECs, live CD45⁺ IELs derived from the epithelial fraction, and live CD45⁺ LPLs from the lamina propria. B cells were depleted using anti-CD19 antibody during FACS to enrich for rare T cell populations. For each sample, post-sort viability was confirmed to be ≥ 90% by trypan blue exclusion, and 10,000 live cells were loaded onto the 10x Chromium Controller. scRNA-seq libraries were prepared using the Chromium Single Cell 3′ Gene Expression Kit (10x Genomics) at the FCSC Core Facility (UAB) according to the manufacturer’s protocol. Libraries from all 52 samples (Supplemental Table) were pooled and sequenced on two Illumina NovaSeq 6000 S4 flow cells using 2 × 100 bp paired-end reads.

### Read alignment and gene quantification

Raw sequencing data were processed using Cell Ranger v9.0.1 (10x Genomics). Raw base call (BCL) files were converted to FASTQ format using cellranger mkfastq with default parameters. Alignment, barcode processing, and UMI counting were performed using the cellranger count pipeline with default settings, aligning reads to the mouse reference genome GRCm39. Gene expression matrices were generated based on valid barcodes identified by Cell Ranger. Data is publicly available at the Gene Expression Omnibus GEO under accession GSE1111111.

### Quality control, normalization, and cell type annotation

All analyses were performed within the scverse ecosystem^15^. Initial preprocessing and quality control were performed using Scanpy (v1.9.8)^16^. Ambient RNA contamination was mitigated using scAR^17^, and doublets were identified and removed using Solo^18^ implemented via the scvi-tools package (v1.1.2)^19^ with default parameters. Cells in which detected genes were < 250 or > 5,500 (±2 SD from the mean) were removed, and genes expressed in fewer than three cells were also excluded. An initial Leiden (v0.10.2)^20^ clustering step separated epithelial (Ep) and immune (Imm) compartments, enabling compartment-specific mitochondrial cutoffs of < 70% for Ep cells and < 25% for Imm cells, each derived from ±2 SD around the mean. Counts were then normalized to 10,000 per cell using scanpy.pp.normalize_total and log-transformed with scanpy.pp.log1p. Further, sc.tl.score_genes_cell_cycle was used to compute S- and G2/M-phase scores for each cell and infer the corresponding cell-cycle phase.

Batch correction and latent embedding were then performed using scVI^21^, incorporating batch information (file_id), continuous covariates (n_genes, total counts, mitochondrial and ribosomal percentages), and relevant categorical covariates (genotype, timepoint, replicate, cell-layer and cell-cycle phase). Post-integration, neighborhood graphs were computed from the scVI latent space, followed by Leiden clustering (resolution = 0.2) and UMAP visualization^22^. Major lineages (T cells, epithelial cells, myeloid cells, B cells, neutrophils, and fibroblasts) were annotated based on canonical marker expression. To resolve finer cell states, each lineage was iteratively subset and re-clustered (resolution 0.3–1.0), to obtain the lowest resolution that yielded biologically distinct, marker-defined subclusters. The sub-clusters were annotated using (a) literature-derived marker genes, (b) the top 250 DEGs per sub-cluster determined using sc.tl.rank_genes_groups, and (c) the anatomical origin of the cells (IEC, IEL, or LPL) (Supplemental Table). In total, 559,621 high-quality cells were retained for downstream analysis. Finally, scANVI^23^ was run to improve biological conservation across genotypes, timepoints, and cell fractions.

### Pseudobulk differential expression testing

Differential expression analysis was performed using a pseudobulk approach with pyDESeq2 (v0.4.12)^24,25^. Cells were grouped by cell state, timepoint and genotype, and raw UMI counts were aggregated to form pseudobulk replicates. DESeq2 normalization, dispersion estimation, and negative binomial generalized linear modeling were performed using the standard workflow, and statistical significance was assessed using the Wald test and p-values were adjusted using the Benjamini–Hochberg FDR method. Genes with an adjusted p-value < 0.05 were considered significantly differentially expressed.

### Gene regulatory network inference with pySCENIC

Transcription factor (TF) regulatory networks were inferred from single-cell RNA-seq data using pySCENIC (v0.12.1)^26,27^ in a lineage-specific workflow. Analyses were performed separately for each major lineage (Epithelial, T, Myeloid, Neutrophil, Innate, B). Within each lineage, genes detected in ≥ 10 cells with ≥ 3 counts were retained, and 8,000 highly variable genes were combined with all detected TFs. To build robust regulatory networks, ∼500 cells per cluster were sampled in a genotype × timepoint-balanced manner across three independent seeds. First, transcription factor–target co-expression modules were identified using GRNBoost2 ^28^, a gradient boosting method that detects genes whose expression patterns covary with each transcription factor across single cells. This step yields candidate TF→target regulatory edges supported by consistent co-expression relationships. Edges reproducibly detected in at least two of the three seeds were retained. Second, motif enrichment pruning was performed using RcisTarget, which evaluates whether candidate target genes contain enriched TF-binding motifs using the mm10 (v10nr) motif databases. Only TF-target pairs with motif support were preserved, and regulons containing ≥ 8 targets were retained. Finally, regulon activity was quantified using AUCell, which computes the Area Under the recovery Curve for each regulon’s gene set within individual cells producing cell-specific regulon activity scores. This score reflects how highly a regulon’s target genes are expressed relative to all other genes in that cell.

Regulon activity differences between the two genotypes were assessed within each cluster by computing, for every regulon, the genotype-specific median AUCell activity and the resulting activity difference (Δ median = *H2-Ab1*^fl/fl^ - *H2-Ab1*^Villin^). For visualization, we generated TF rewiring bar plots, showing the top regulons enriched per cluster in both genotypes based on |Δ median|, with positive values indicating higher activity in *H2-Ab1*^fl/fl^ and negative values indicating higher activity in *H2-Ab1*^Villin^.

### Ligand-receptor interaction analysis

Cell-cell communication was inferred using LIANA+ (v1.5.1)^28,29^, employing the rank_aggregate method to score ligand–receptor interactions. A unified ligand–receptor reference was constructed by concatenating multiple curated resources (Consensus, MouseConsensus, Baccin2019, CellChatDB, CellPhoneDB, ConnectomeDB2020, Embrace, Ramilowski2015, ICELLNET, and LRdb) and removing duplicates. LIANA was run per tissue layer (epithelial or lamina propria) and timepoint, using annotated cell states as sender and receiver identities, with communication scores computed per sample, where each sample was defined by tissue layer × timepoint × genotype × replicate. LIANA’s rank_aggregate.by_sample method was used to compute interaction scores per sample, retaining only ligand–receptor pairs with sufficient biological support—at least 5 cells per sender–receiver group and expression in ≥ 5% of cells expressing the ligand or the receptor.

Differential ligand–receptor activity between *H2-Ab1*^fl/fl^ and *H2-Ab1*^Villin^ was assessed using a custom LIANA+ workflow. Per-sample LIANA scores were aggregated across replicates to compute genotype-specific medians for three metrics: (i) *mag_score* (–magnitude_rank, where higher values indicate stronger interactions), (ii) *lr_means* (mean ligand and receptor expression within sender and receiver), and (iii) *expr_prod* (ligand × receptor expression product). Only interactions with ≥ 2 biological replicates per genotype were retained. Differential effects for the above three parameters were calculated as (*H2-Ab1*^fl/fl^ - *H2-Ab1*^Villin^). The resulting effect-size measures (*mag_score_effect*, *lr_means_effect*, and *expr_prod_effect*) were used to identify sender→receiver interactions enriched in either genotype, and only interactions with absolute magnitude of differential effect (|*mag_score_effect*|) ≥ 0.15 were retained. Visualization: For each receiver subset, the top ten ligand–receptor interactions per sender, ranked by |*mag_score_effect*| were selected, and differential communication patterns were visualized using scatterplots of *lr_means_effect* versus *expr_prod_effect*.

## Extended Data Figures

**ED Fig. 1:**
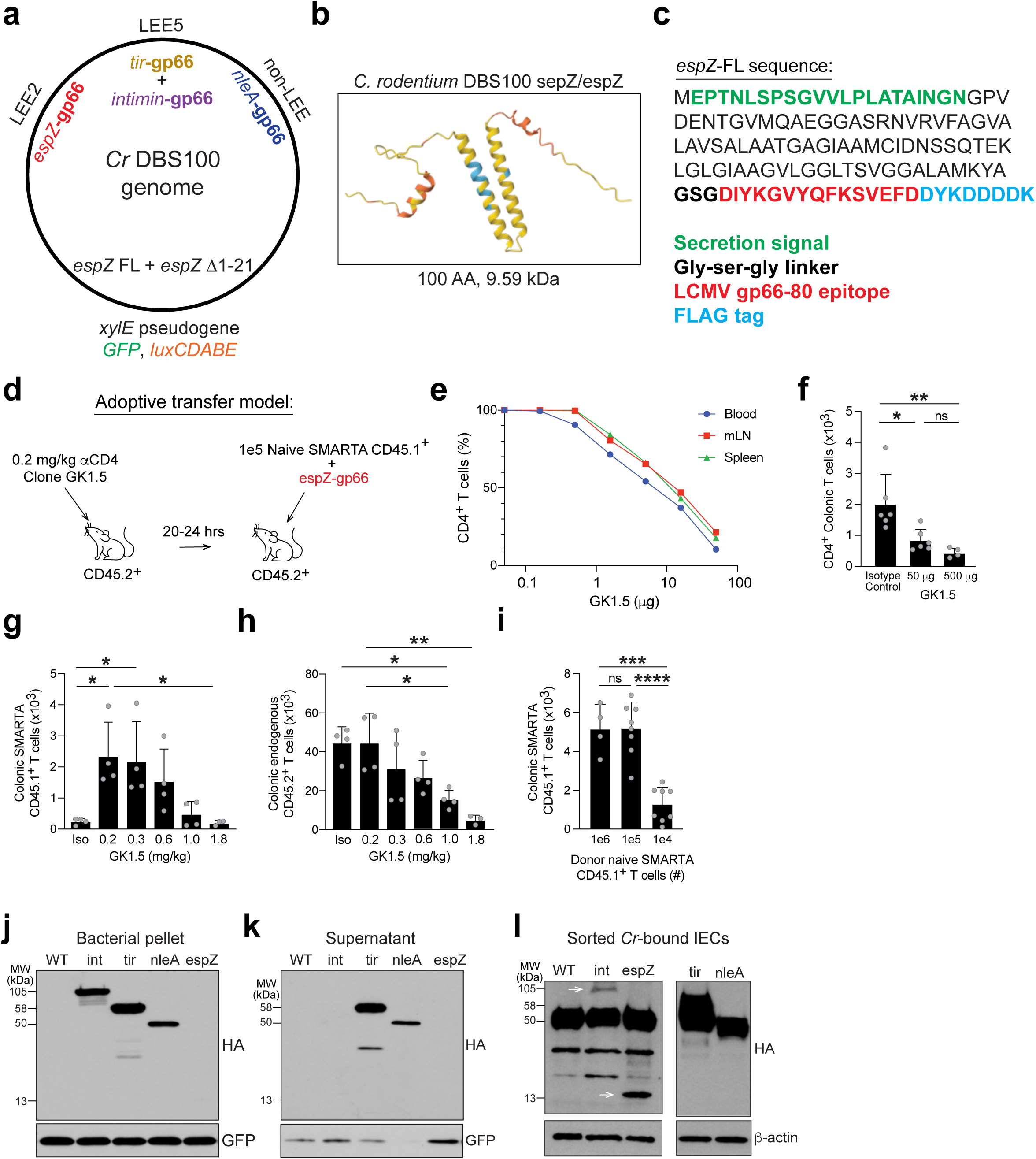
Generation of epitope-tagged *C. rodentium* strains and optimization of adoptive transfer model. **a**, Schematic of *Cr* DBS100 genome with LEE regions annotated for intimin (*eae*), *tir*, and *espZ* in DBS100 parent strain; non-LEE region annotated for *nleA* in DBS100 parent strain; and *xylE* pseudogene annotated for insertion of full-length espZ-gp66 and espZ D1-21-gp66 replacing coding sequence for GFP in parent *Cr*-GFP strain. **b**, 3D ribbon diagram of *C. rodentium* strain DBS100 sepZ/espZ protein generated with AlphaFold3 with per-residue confidence metric (pIDDT) very high (pIDDT>90; dark blue), confident (90>pIDDT>70; cyan), low (70>pIDTT>50; yellow), and very low (pIDDT<50; orange). **c**, Full-length espZ protein sequence with T3SS secretion signal (green), linker (black), LCMV gp66-80 epitope sequence (red), and FLAG tag (blue) annotated. **d**, Adoptive transfer model used throughout this study whereby recipient C57BL/6 CD45.2^+^ recipient mice are administered 0.2 mg/kg anti-CD4 antibody 20-24 hours prior to receiving 1e5 naïve CD45.1^+^ SMARTA T cells and infection with espZ-gp66 (red) to create niche space in T-cell repertoire for engrafted SMARTA cells. **e**, Optimization of anti-CD4 (GK1.5) antibody administration across 0.1-10 mg doses on recovered CD4^+^ T cells from blood (blue), mesenteric lymph node (mLN; red), or spleen (green). **f**, Number of colonic CD4^+^ T cells recovered after depletion with anti-CD4 antibody or isotype control. 3-5 mice per group; *n*=2 independent experiments. One-way ANOVA; *p≤0.05 and **p≤0.01. **g,h**, Number of colonic CD45.1^+^ SMARTA T cells (**g**) and endogenous CD45.2^+^ T cells (**h**) recovered from mice depleted with 0.2-1.8 mg/kg anti-CD4 or isotype control 14 days after adoptive transfer and infection with espZ-gp66. 3-5 mice per group; *n*=2 independent experiments. One-way ANOVA; *p≤0.05 and **p≤0.01. **i**, Number of colonic CD45.1^+^ SMARTA T cells recovered from mice depleted with 0.2 mg/kg anti-CD4 14 days after adoptive transfer with 1e4-1e6 naïve CD45.1^+^ SMARTA cells and infection with espZ-gp66. 3-5 mice per group; *n*=2 independent experiments. One-way ANOVA; ***p≤0.001 and ****p≤0.0001. Western blot for HA tag and GFP loading control performed on *Cr*-GFP parent strain and intimin-gp66, tir-HA, nleA-gp66, and espZ-gp66 *Cr*-GFP derivative bacterial cell pellets (**j**) and bacterial supernatants (**k**) from culture in DMEM. **l**, Western blot for HA tag and β-actin loading control performed on sorted *Cr*-bound IECs infected with *Cr*-GFP parent strain and intimin-gp66, tir-HA, nleA-gp66, and espZ-gp66 *Cr*-GFP derivatives.

**ED Fig. 2:**
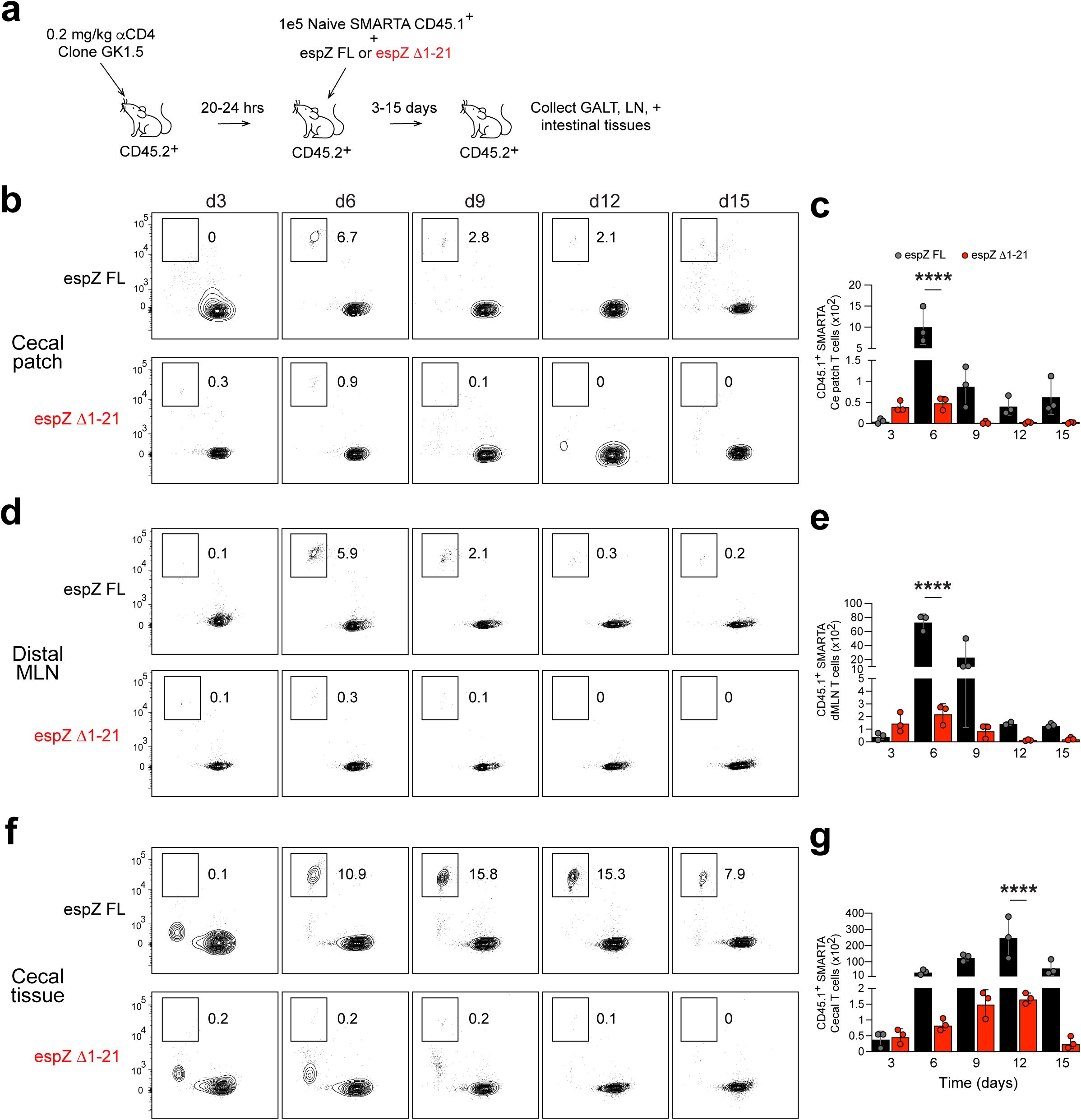
Kinetics and tissue distribution of SMARTA T cell responses to translocated versus bacterially-associated espZ antigens. C57BL/6 CD45.2^+^ mice received 0.2 mg/kg anti-CD4 20-24 hours prior to adoptive transfer of 1×10^5^ naïve CD45.1^+^ SMARTA T cells and infection with either espZ FL-gp66 or espZ D1-21-gp66 strains. **a**, Recovered colonic CD4^+^TCRβ^+^ cells were analyzed by flow cytometry for CD45.1 or CD45.2 expression across GALT (cecal patch and distal mesenteric LN) and cecal tissue at days 3-15 post-infection. **b-g**, Number of recovered CD45.1^+^ SMARTA T cells from cecal patch (**b,c**), distal MLN (**d,e**), or cecal tissue (**f,g**). 3-5 mice per group; *n*=2 independent experiments. One-way ANOVA; *p≤0.05, **p≤0.01, and ***p≤0.001.

**ED Fig. 3:**
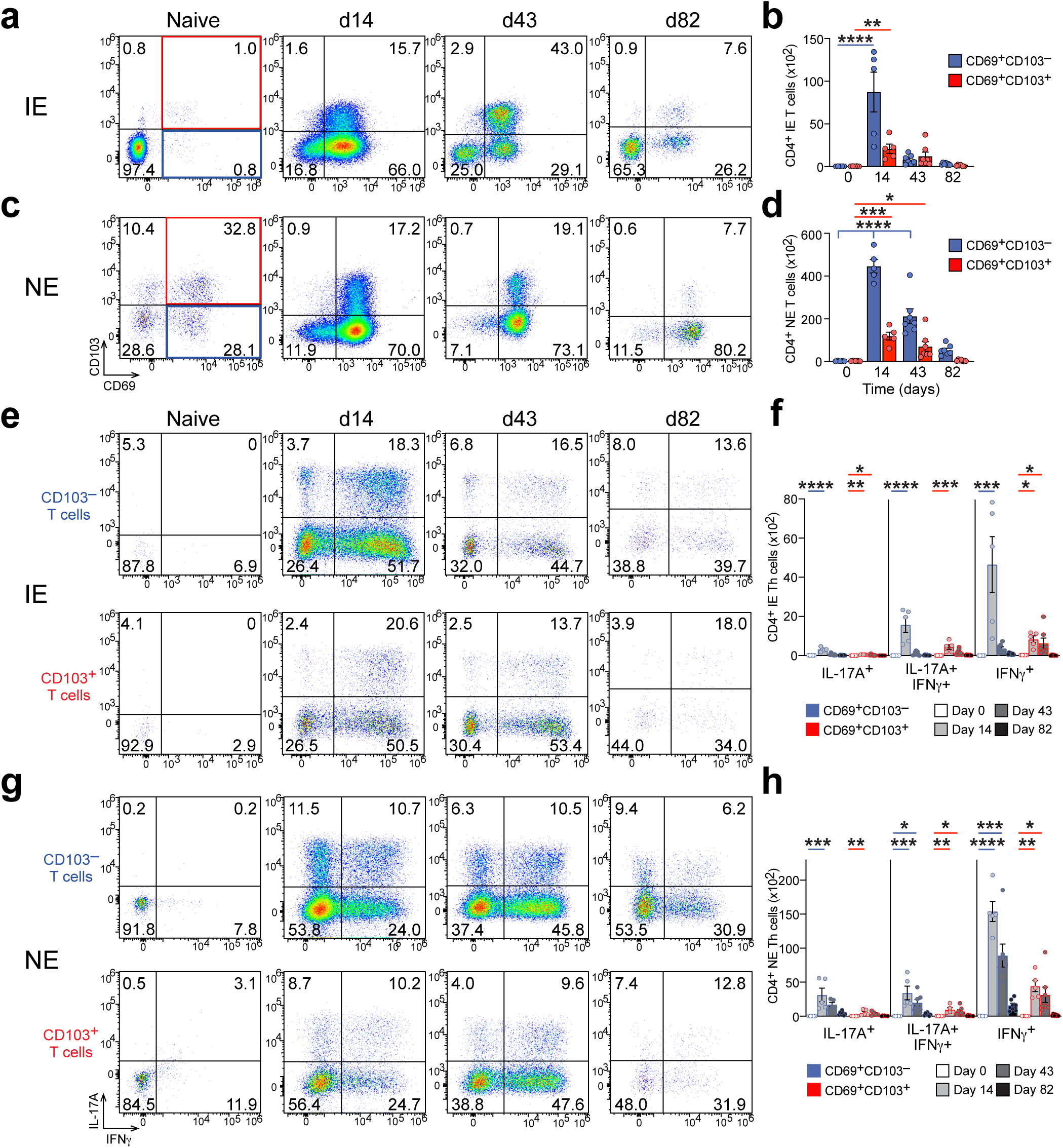
Phenotypic and functional characterization of endogenous CD4^+^ T cells during *C. rodentium* infection. a-h, Colonic CD4^+^TCRβ^+^ cells isolated from colonic epithelium (IE) and non-epithelial tissue (NE) were collected from naïve or *Cr*-DBS100-infected C57BL/6 mice at day 14, 43, or 82 post-infection. Recovered cells were stimulated for 3-4 hours with PMA and ionomycin, stained for surface CD4, TCRβ, CD69, and CD103 followed by intracellular staining for IL-17A and IFNγ and analyzed by flow cytometry. 5-7 mice per group; *n*=2 independent experiments. One-way ANOVA (comparing CD69^+^CD103^−^ populations (blue); comparing CD69^+^CD103^+^ populations (red)); *p≤0.05, **p≤0.01, ***p≤0.001, and ****p≤0.0001.

**ED Fig. 4:**
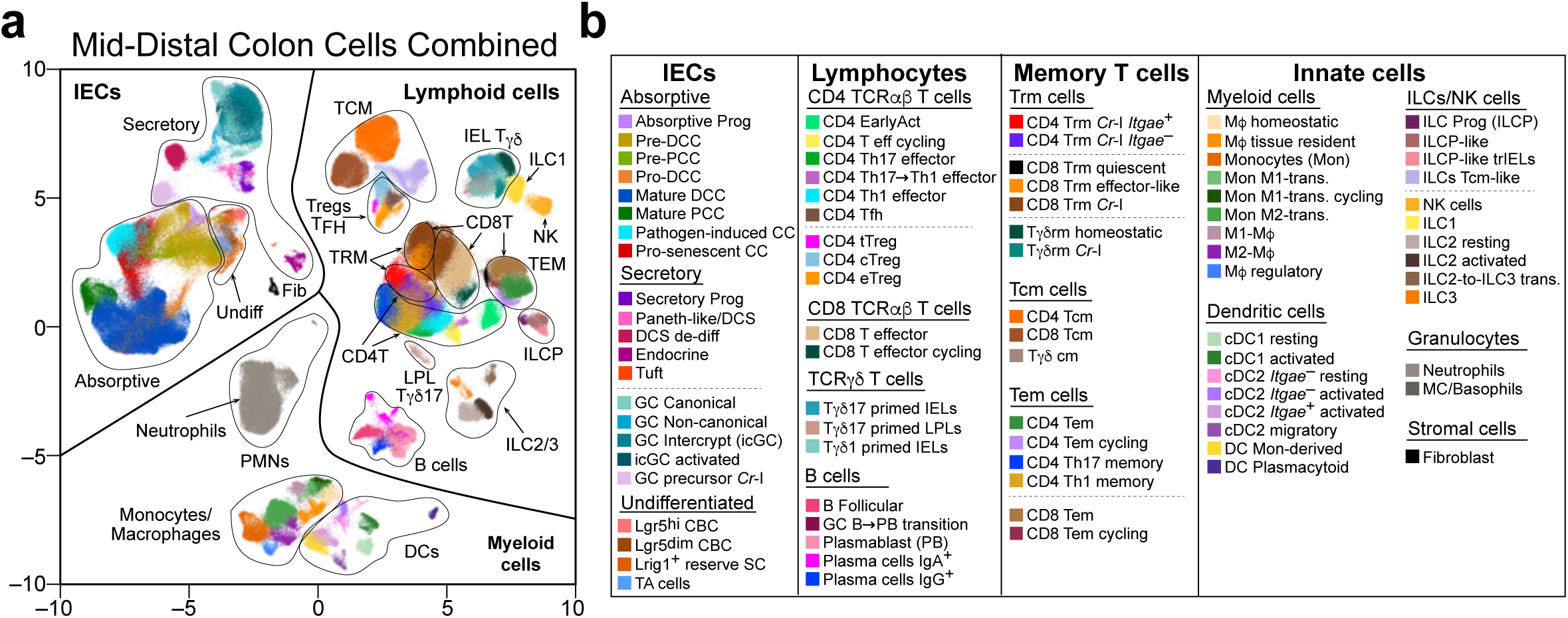
Composite single-cell transcriptomic atlas of colonic immune and epithelial cells from naive and *Cr*-infected *H2-Ab1*^fl/fl^ and *H2-Ab1*^Villin^ mice. a,b, UMAP analysis of total mid-distal colon cells isolated from naïve or DBS100 *Cr*-infected *H2-Ab1*^fl/fl^ or *H2-Ab1*^Villin^ mice on day 9, 14 or 42 post-infection. EpCAM^+^ and CD45^+^ intraepithelial (IE) and non-epithelial (NE) cells were sorted for scRNA-seq analysis. Approximately 600,000 cells were resolved into 87 individual clusters: 22 IEC clusters (8 absorptive lineage, 10 secretory lineage, and 4 undifferentiated populations); 35 lymphocyte clusters (19 effector and 16 memory populations spanning CD4, CD8, *gd* T cells, and B cells); and 30 innate cell populations spanning monocytes/macrophages, dendritic cells, ILCs/NK cells, granulocytes, and stromal fibroblasts. Abbreviations: prog, progenitor, DCC, distal colonocyte; PCC, proximal colonocyte; CC, colonocyte; DCS, deep crypt secretor; de-diff, de-differentiated; GC, goblet cell; *Cr*-I, *C. rodentium*-induced; CBC, crypt base columnar; TA, transit amplifying; EarlyAct, early activated; T eff, T effector; Tfh, T follicular helper; tTreg, thymic Treg; cTreg, central Treg; eTreg, effector Treg; GC B, germinal center B cell; PB, plasmablast; Trm, tissue resident memory T cell; Tcm, central memory T cell; Tem, effector memory T cell; ILCP, innate lymphoid cell precursor; trIELs, tissue resident intraepithelial lymphocytes; NK, natural killer; MC, mast cell.

**ED Fig. 5:**
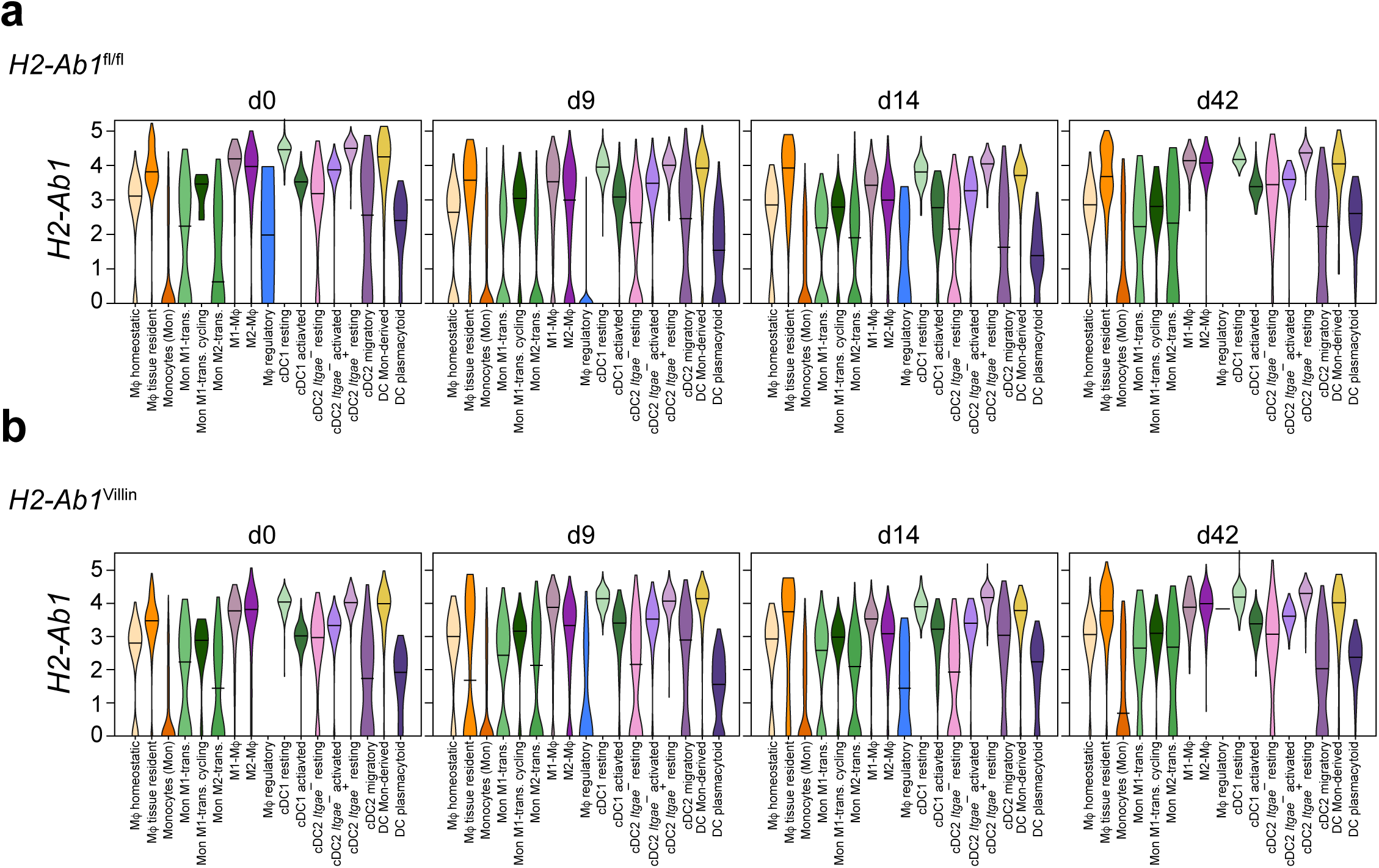
Temporal expression of *H2-Ab1* across professional antigen-presenting cells in *H2-Ab1*^fl/fl^ and *H2-Ab1*^Villin^ mice. **a-b**, Violin plots of *H2-Ab1* expression across monocyte/macrophage and dendritic cell clusters in naïve or DBS100 *Cr*-infected *H2-Ab1*^fl/fl^ or *H2-Ab1*^Villin^ mice on day 9, 14 or 42 post-infection. Abbreviations: DC, dendritic cell; Mf, macrophage; trans, transitioning; Mon, monocyte.

**ED Fig. 6:**
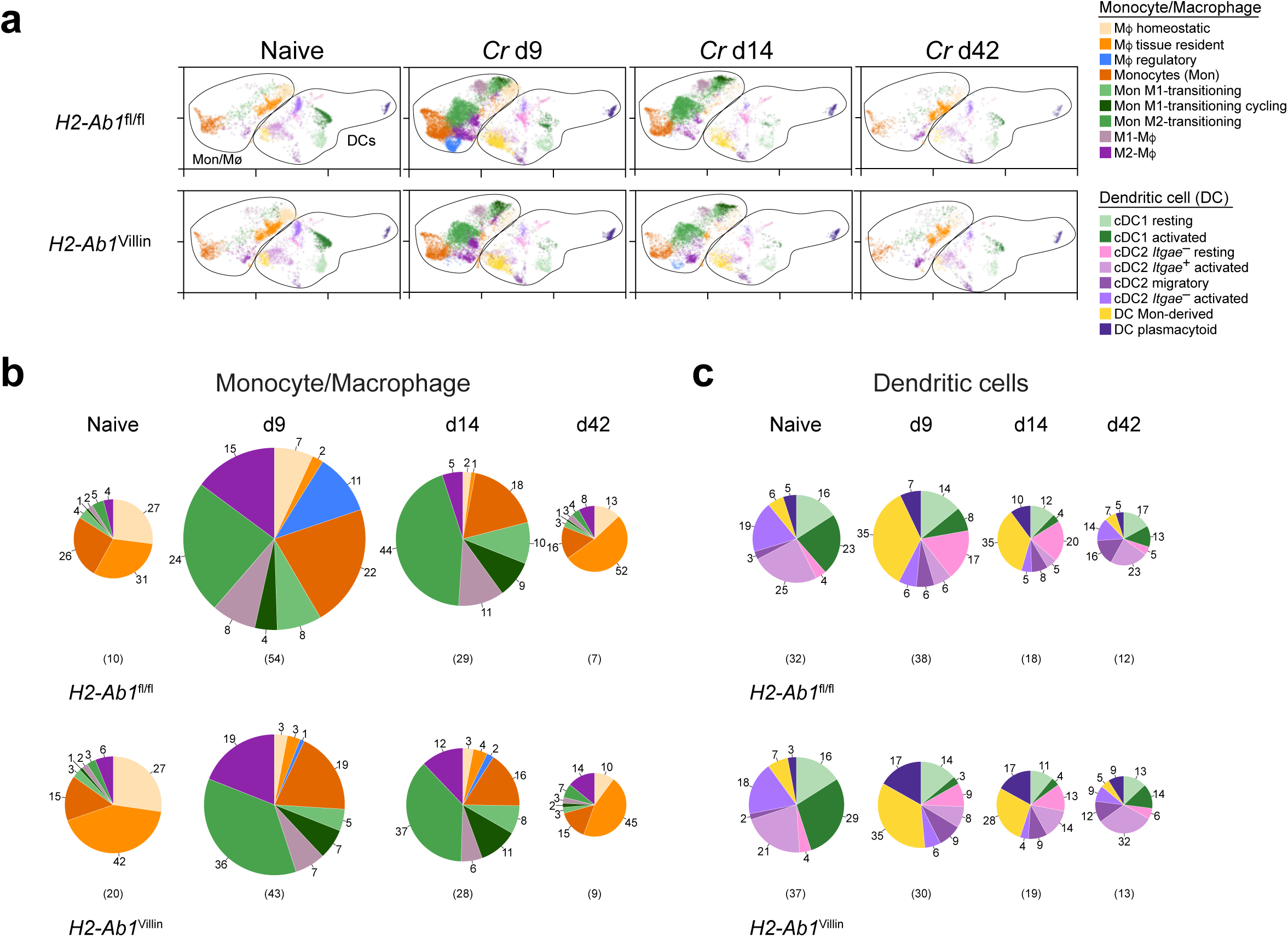
Distribution of myeloid cell populations across intraepithelial and non-epithelial compartments in *H2-Ab1*^fl/fl^ and *H2-Ab1*^Villin^ mice. **a**, UMAPs of total monocyte/macrophage and dendritic cell populations isolated from mid-distal colon of naïve or DBS100 *Cr*-infected *H2-Ab1*^fl/fl^ or *H2-Ab1*^Villin^ mice on day 9, 14 or 42 post-infection. **b**,**c,** Pie charts of the number and percentages of monocytes/macrophages (**b**) and dendritic cells (**c**) at indicated times, highlighting that while total monocyte/macrophage numbers and subset composition are substantially altered within and between genotypes during active infection (d9), total DCsand DC subsets do not undergo major shifts in numbers or subset composition between *H2-Ab1*^fl/fl^ and *H2-Ab1*^Villin^ mice. Pie chart areas normalized to largest population within monocyte/macrophage or DC pools.

**ED Fig. 7:**
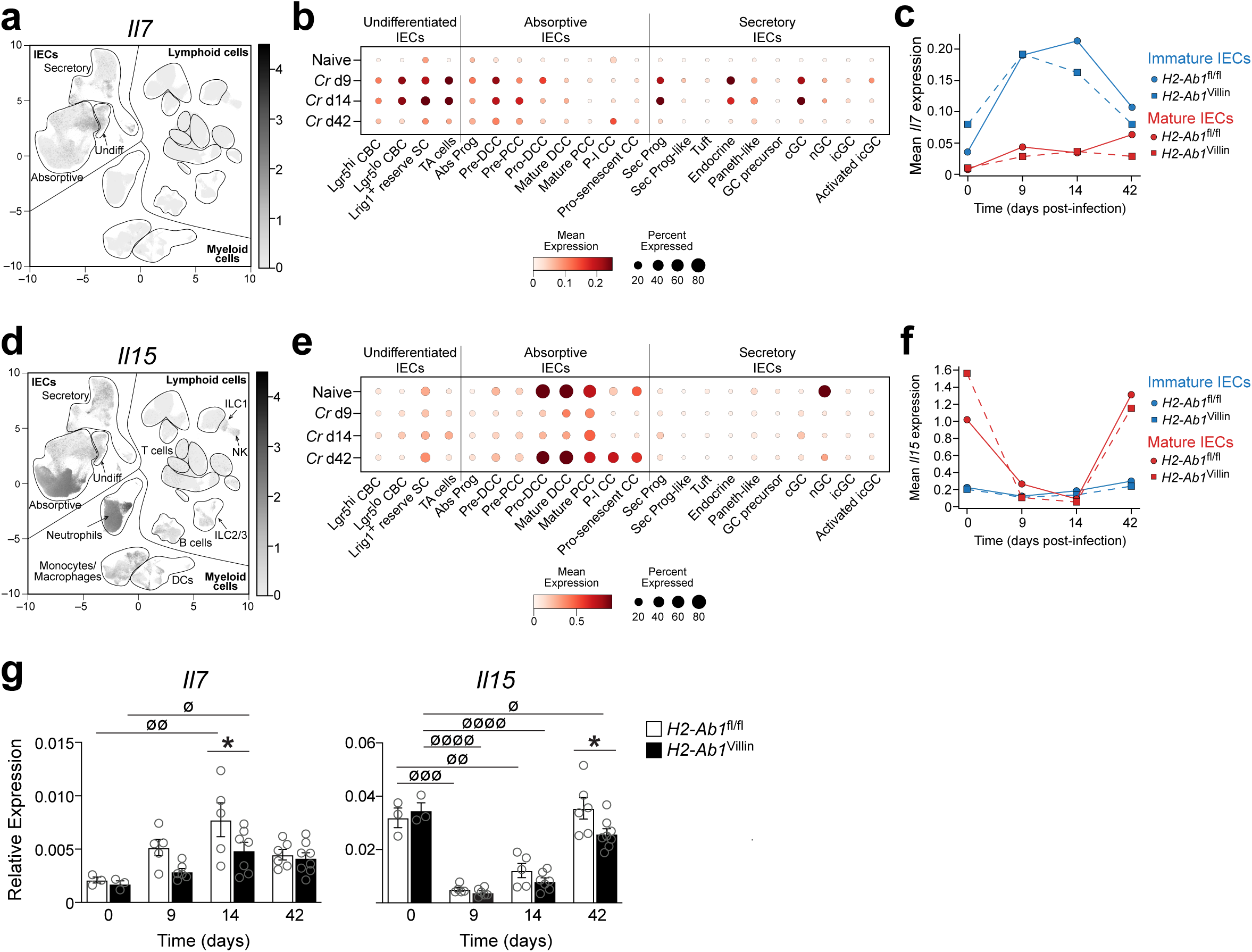
Epithelial *Il7* and *Il15* expression shows reciprocal temporal regulation and cellular source during *Cr* infection. **a**, UMAP analysis of *Il7* expression in scRNA-seq dataset. **b**, Dot plot of *Il7* expression across time points and cell types of *H2-Ab1*^fl/fl^ mice during *Cr* infection, highlighting significant *Il7* expression by undifferentiated IECs (Lgr5dim CBCs, Lrig1^+^ reserve SCs, and TA cells), immature absorptive IECs (pre-DCCs and pre-PCCs), and immature secretory IECs (secretory progenitors), along with endocrine and canonical GCs, during active infection (days 9-14). **c**, Comparison of expression of *Il7* by immature and mature IECs of *H2-Ab1*^fl/fl^ and *H2-Ab1*^Villin^ mice over infection time course, highlighting similar pattern of expression by immature IECs of both genotypes that is maximal during active infection (days 9 and 14). **d**, UMAP analysis of *Il15* expression in scRNA-seq dataset. **e**, Dot plot of *Il15* expression across time points and cell types of *H2-Ab1*^fl/fl^ mice during *Cr* infection, highlighting significant *Il15* expression by mature absorptive IECs (pro-DCC, mature DCC, mature PCC, P-I CC, and pro-senescent CC) as well as some non-canonical GCs at homeostatic time points (naive and day 42) but diminished expression during acute infection (days 9-14)—displaying a reciprocal expression pattern from that of *Il7*. **f**, Comparison of expression of *Il15* by immature and mature IECs of *H2-Ab1*^fl/fl^ and *H2-Ab1*^Villin^ mice over infection time course, highlighting enhanced expression of *Il15* in mature IECs compared to immature IECs at homeostatic time points and similar pattern of expression between genotypes. **g**, *Il7* and *Il15* expression from sorted cECs isolated from mid-distal colon of naïve or DBS100 *Cr*-infected *H2-Ab1*^fl/fl^ or *H2-Ab1*^Villin^ mice on day 9, 14 or 42 post-infection was detected by RT-PCR, confirming the pattern shown in the scRNA-seq dataset. 5-8 mice per group; *n*=2 independent experiments. Two-way ANOVA; *p≤0.05 (comparing *H2-Ab1*^fl/fl^ *H2-Ab1*^Villin^ mice); ^∅^p≤0.05, ^∅∅^p≤0.01, ^∅∅∅^p≤0.001, and ^∅∅∅∅^p≤0.0001 (comparing timepoints). IN view of the fact that IL-7 and IL-15 are key common γ-chain cytokines that support survival of different Trm subsets, our data indicate that there are temporal and spatial expression differences between the two cytokines that emerge from the scRNA-seq dataset: *Il7* transcripts were predominantly expressed by immature, undifferentiated cECs (CBCs, TA cells, pre-DCCs, and some secretory cells) at days 9-14 post-infection, whereas *Il15* transcripts were predominantly expressed by mature IECs (pro-DCCs, mature DCCs, and pro-senescent CCs) in naïve cECs and at day 42 post-infection. The loss of *Il15* expression at days 9 and 14 likely reflected more rapid epithelial turnover and alterations in DCC maturation that accompany active *Cr* infection, as mature cells at mucosal surface are rapidly shed and replaced by immature populations. These findings suggest that IL-15 plays a key role in maintenance of intraepithelial T cells at homeostasis, and when the cellular source of IL-15 is lost during acute infection, IL-7 from proliferating immature cells—and possibly IL-2 produced by activated T effector cells—may compensate for this loss as new memory cells are generated and occupy the epithelial niche.

